# mTOR signaling promotes cytokine production in T cells through 3’UTR-mediated translation control

**DOI:** 10.1101/2024.10.01.616022

**Authors:** Anouk P. Jurgens, Josephine Zwijnen, Floris P.J. van Alphen, Antonia Bradarić, Koos Rooijers, Arie J. Hoogendijk, Branka Popović, Monika C. Wolkers

## Abstract

T cells are key contributors to clear our body from infected and malignant cells. When T cells respond to target cells, they undergo profound translational alterations. The evolutionary and highly conserved kinase mammalian target of rapamycin (mTOR) is a central mediator of T cell differentiation, homeostasis, and T cell activation, including the production of the key pro-inflammatory cytokines TNF, IL2, and IFNγ. mTOR was shown to execute its translation activity through TOP motifs located in the 5’ Untranslated region (5’UTR) of its target genes. Here, we uncovered a distinct mechanism of mTOR signaling on cytokine production in T cells, which is under control of the 3’UTR. Even though non-classical TOP motifs are present in cytokine 3’UTRs, they do not contribute to mTOR-mediated translation regulation. Rather, AU-rich elements (AREs) are required for mTOR-mediated cytokine production. Furthermore, we discovered that the RNA binding protein DDX21 binds to 3’UTR AREs and confers the mTOR-mediated translation control. In conclusion, we here present a previously unappreciated ARE-dependent, 3’UTR-mediated mode of action that mTOR employs to regulate cytokine production.

## INTRODUCTION

T cells constitute a critical component of the adaptive immune system against pathogens and cancer cells. To perform their function effectively, T cells require precise control of protein expression, which is in particular important during T cell activation, when they produce key effector molecules such as pro-inflammatory cytokines ^1,2^. mTOR is one of the key regulators of T cell biology, as its signaling promotes T cell growth, proliferation and effector function, and regulates T cell fate decisions ^3–9^. Likewise, mTOR signaling regulates the rate of protein translation by integrating the signals a T cell receives through the T cell receptor (TCR), co-stimulatory receptors, and cytokine receptors ^7,10–13^. mTOR-signaling also coordinates the fine-tuning of gene expression ^5^. For instance, mTORC1 regulates the cytotoxicity in murine T cells, and its inhibition hampers the expression of various effector molecules such as IFNγ ^6^. However, how mTOR signaling acts on its target mRNAs, and how it promotes cytokine expression in T cells is, to date not fully understood.

In addition to the coding sequence, sequences within the untranslated regions (UTRs) of the mRNA, i.e. the 5’UTR and 3’UTR, are key contributors to translation control. In particular RNA-binding proteins (RBPs) convey the regulation of - amongst other processes - RNA splicing, poly-adenylation, localization, stability and translation of (m)RNA ^2,14–19^. mTOR-mediated regulation of translation can also be mediated by RBPs: mTOR phosphorylates the translation inhibitor LARP, resulting in its dissociation from cytidine and pyrimidine-rich 5’terminal oligopyrimidine (TOP)-like motifs in the 5’UTR of its target mRNAs ^20–22^. The release of LARP from its target mRNA, along with the mTOR mediated phosphorylation of the translation inhibitor 4E-BP allows the eukaryotic initiation factors (eIFs) to engage with the 5’UTR and to initiate translation ^23^. However, also the 3’UTR can contribute to translation control ^18^, and mTOR signaling could possibly contribute to this regulation. In mouse embryonic fibroblasts, mTOR signaling was reported to modulate translation by 3’UTR shortening, which led to increased polysome association and translation of its target genes ^24^. Whether mTOR signaling can also directly contribute to the 3’UTR-mediated translation control in T cells has not been studied yet is conceivable provided its key role in regulating protein expression.

In this study, we uncovered that mTOR signaling coordinates the translation of the three key pro-inflammatory cytokines TNF, IL2 and IFNγ in human T cells through their 3’UTR. Surprisingly, we found that the non-classical TOP-like motifs present in all three cytokine 3’UTRs are redundant. Rather, AU-rich elements (ARE) are required for the mTOR-mediated translation control. With RBP-pulldown assays, we identified several RBPs that engage with AREs in the IL2 3’UTR in an mTOR-dependent manner. Specifically, DDX21 is required for the IL2 3’UTR specific regulatory function of mTOR. Combined, we reveal that mTOR-mediated regulation of ARE binding proteins to 3’UTRs of target mRNAs supports the protein production upon T cell activation.

## RESULTS

### mTOR negatively influences cytokine production in human T cells

To study the mTOR-mediated regulation of cytokine production in effector CD3^+^ T cells (Teff), we activated human peripheral blood-derived T cells for 48h with αCD3/αCD28, which were then rested for 96h. When reactivated with phorbol myristate acetate (PMA)/ionomycin, T cells display similar rapid cytokine production kinetics as *ex vivo* activated Teff and memory T cells ^25–27^. Blocking mTOR during T cell activation with the ATP-competitive mTORC1/mTORC2 inhibitor Torin-1 (Supplemental Figure 1A) ^4,6,28,29^, or with the mTORC1 selective, allosteric, inhibitor Rapamycin ^4,6,29^ did not affect cell viability, nor the induction of the early T cell activation marker CD69 when compared to DMSO control-treated T cells (Supplemental Figure 1 B,C). Yet, and as previously reported for murine T cells ^6^, mTOR inhibition by Torin-1 significantly decreased the production and secretion of TNF and IL2 and showed a trend for IFNγ in human T cells (Supplemental Figure 1D, E). Similar effects were observed on cytokine production upon Rapamycin treatment (Supplemental Figure 1F), indicating that the cytokine production in T cells is primarily regulated through mTORC1. Notably, whereas mTOR inhibition affected the cytokine production, effects on cytokine mRNA levels and RNA stability were limited (Supplemental Figure 1G). Thus, mTOR signaling promotes protein production of pro-inflammatory cytokines in human T cells, and it does so through translation control.

### mTOR-mediated regulation of protein expression employs cytokine 3’UTRs

Next, we questioned how mTOR regulates the production of cytokines in human CD8^+^ T cells. We and others previously reported the importance of cytokine 3’UTRs in fine-tuning protein production ^14,16–18,30,31^. This became particularly evident with full length *TNF*, *IL2,* or *IFNG* 3’UTR GFP reporter constructs that substantially increased their protein expression upon T cell activation^31^. Whether mTOR signaling contributes to the 3’UTR-mediated protein regulation in T cells, has however not yet been studied. To test this, we measured the effect of Torin-1 treatment on *TNF*, *IL2,* or *IFNG* 3’UTR GFP reporter constructs. *GZMB* 3’UTR that lacks regulatory capacities in activated T cells^31^ and GFP fused to no 3’UTR (GFP-control) served as control.

Torin-1 treatment did not affect GFP reporter expression in resting T cells (Figure 1A & Supplemental Figure 2A). Likewise, when Teff cells carrying GFP-control or GFP-*GZMB* 3’UTR were activated with PMA/Ionomycin, only mild effects on GFP protein expression levels were observed when mTOR signaling was blocked (Figure 1A & Supplemental Figure 2A). In sharp contrast, the induction of GFP protein expression under control of the three cytokine 3’UTRs was substantially impaired when mTOR was blocked with Torin-1 or Rapamycin, a feature that was most prominent for the *IL2* and *IFNG* 3’UTR (Figure 1A & Supplemental Figure 2B). Similar results were obtained upon αCD3/αCD28 stimulation (Supplemental Figure 2C). Of note, this mTOR-sensitive, 3’UTR-mediated regulation of protein expression was conserved in human CD4^+^ T cells (Supplemental Figure 2D). In conclusion, mTOR signals through cytokine 3’UTRs to promote protein production in T cells.

**Figure 1.**
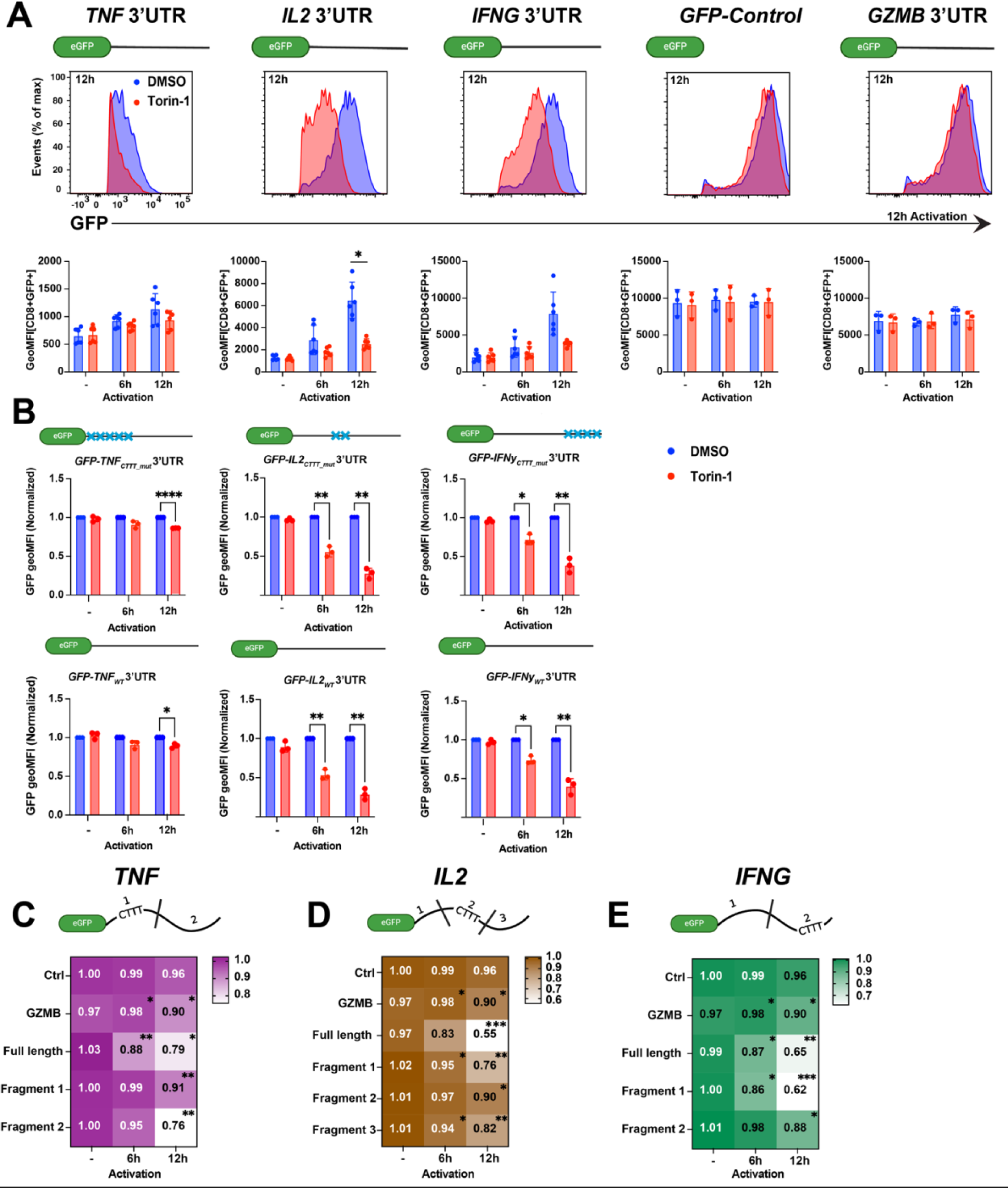
mTOR boosts cytokine production through the cytokine 3’UTR independently of TOP-like motifs. A) Human CD8^+^ Teff cells expressing indicated full-length 3’UTR fused to a GFP reporter, or GFP empty control (*GFP-Control*) were activated for indicated time points with PMA/Ionomycin in the presence of 250nM Torin-1 (red) or 0.05% DMSO control (blue). Top panel: Representative GFP expression at 12h post activation as determined by flow cytometry. Bottom panel: Compiled GFP geoMFI from two independently performed experiments. (-) indicates non-activated T cells treated with Torin-1 or DMSO for 12h. (n=3-6 donors, mean ± SD) B) Teff cells expressing GFP with indicated WT cytokine 3’UTRs or CTTT mutant variants thereof (CTTTàCTGT) were activated with PMA/Ionomycin in the presence of Torin-1 or DMSO. Each GFP geoMFI expression was normalized to its paired DMSO control. (n=3 donors, mean ± SD). C) -E) Teffs were transduced with full length *TNF* (C), *IL2* (D) and *IFNG* (E) 3’UTRs, or with indicated fragments (fragments containing CTTT motifs indicated) and activated with PMA/Ionomycin in the presence of Torin-1 or DMSO. GFP *Control* and *GZMB* served as control. Heatmap scale representative of GFP ratio between DMSO/Torin-1 treated cells. (n=3 donors). A-C: One sided paired student t test; *p < 0.05, **p < 0.01, ***p < 0.001. Also see Supplemental Figure 1, 2.

### TOP-like motifs in the 3’UTR do not contribute to mTOR mediated protein production

We then sought to define which motifs in the cytokine 3’UTRs mediate the mTOR-driven regulation. mTOR signaling can phosphorylate the translation inhibitor LARP1, which then dissociates from CTTT-rich TOP-like motifs in the 5’UTR of its target genes ^21,23,32^. Interestingly, LARP1 was recently shown to interact with 3’UTRs in HEK293T cells ^21^. Therefore, we searched for TOP-like CTTT motifs in the cytokine 3’UTRs, and we found them present in all three cytokine 3’UTRs (Supplemental Figure 2E). To test whether the TOP-like motifs were sensitive to mTOR signaling, we mutated the CTTT motifs into CGTT in the full length 3’UTRs (Figure 1B). Surprisingly, these mutants were equally responsive to Torin-1 as the WT cytokine 3’UTRs (Figure 1B), indicating that TOP-like CTTT motifs in the 3’UTRs did not contribute to mTOR-mediated protein expression. When re-examining the cytokine 3’UTRs, we observed that the CTTT motifs clustered together (Supplemental Figure 2E). We therefore generated CTTT-rich and CTTT-poor fragments for each cytokine 3’UTR to identify the Torin-1 sensitive fragment. Again, GFP reporters containing the CTTT-rich fragments poorly responded to Torin-1 treatment (Figure 1C-E). Conversely, the CTTT-poor fragments of all three cytokine 3’UTRs were sensitive to Torin-1 treatment and displayed significantly reduced protein expression upon T cell activation (Figure 1C-E). In conclusion, TOP-like CTTT motifs in cytokine 3’UTRs do not contribute to mTOR-mediated regulation of protein production.

### mTOR regulates 3’UTR-mediated protein expression through AU-rich elements

To uncover which motifs in the cytokine 3’UTRs were responsive to mTOR signaling, we zoomed in on the Torin-1 sensitive 3’UTR fragments. The CTTT-poor regions of all three cytokine 3’UTRs contained AU-rich elements (AREs), sequences that were identified as key hubs for RBPs for regulating cytokine expression (Supplemental Figure 2E) ^15,17,18,33–35^. To study the contribution of AREs to the mTOR-mediated regulation of protein expression, we mutated the AREs, defined here as TATTTA, to TATGTA in the full-length cytokine 3’UTRs (Supplemental Figure 2E). Strikingly, at all time points measured, mutating AREs in the cytokine 3’UTRs substantially reduced the effect of mTOR inhibition on protein expression (Figure 2A-C). The effect of ARE mutations on protein expression was detected for all cytokine 3’UTRs but was most prominent for the *IL2* and *IFNG* 3’UTRs (Figure 2A-C). Thus, ARE sequences in cytokine 3’UTRs mediate the mTOR signaling effects on protein production.

**Figure 2.**
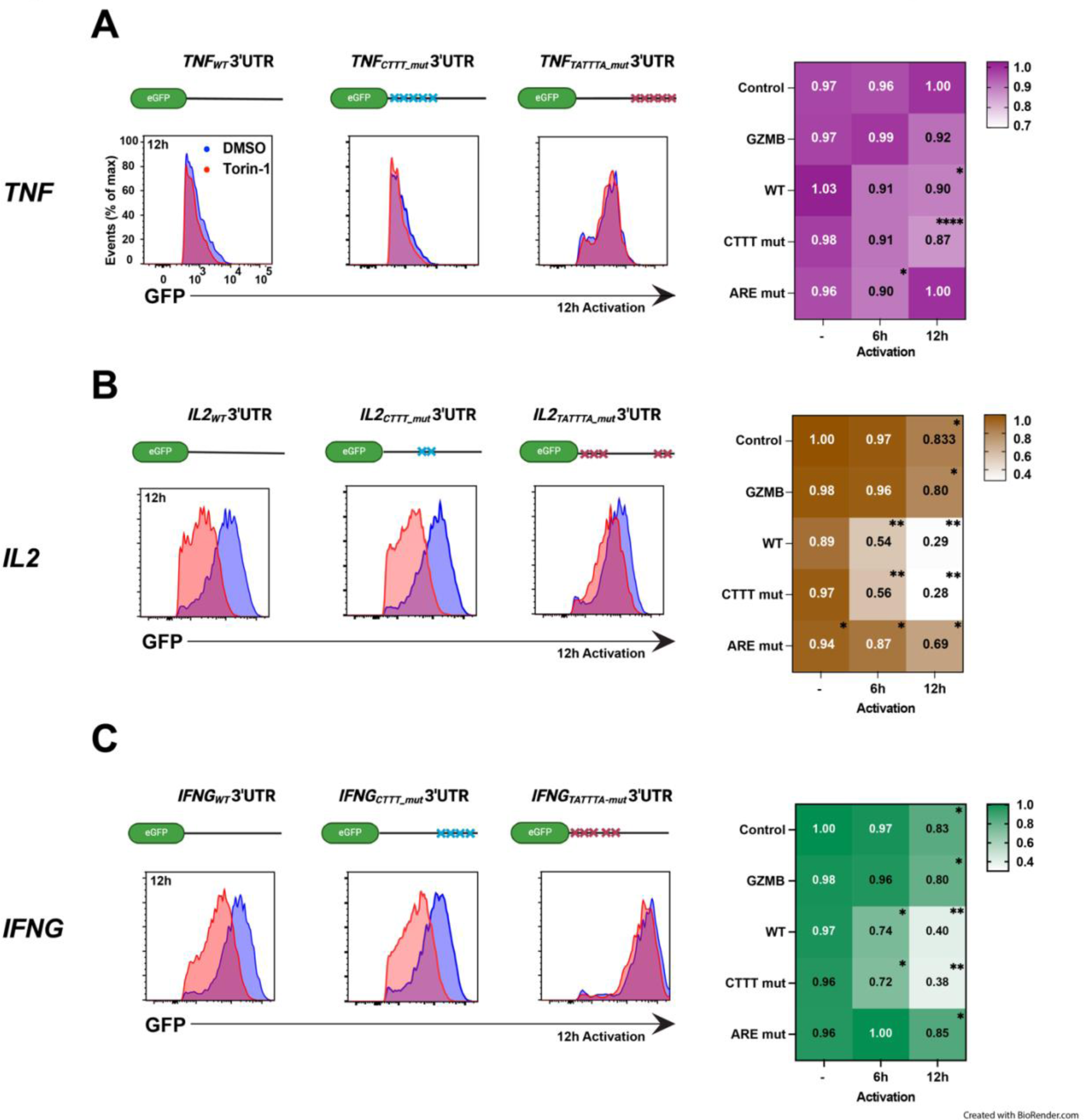
mTOR promotes 3’UTR-mediated protein expression through AU-rich elements. A) -C) CD8^+^ Teff cells expressing indicated full-length *TNF* (A), *IL2* (B) and *IFNG* (C) 3’UTRs or the respective CTTT mutants (CTTT -> CGTT) or ARE mutants (TATTTA -> TATGTA), *Control* or *GZMB* 3’UTR GFP controls were activated with PMA/Ionomycin in the presence of Torin-1 (red) or DMSO (blue). Left: Representative GFP expression at 12h post activation. Right: Heatmap depicting the GFP ratio between DMSO/Torin-1 treated cells. (n=3 donors). (-) indicates non-activated T cells that were treated with Torin-1 or DMSO for 12h. Data of WT cytokine 3’UTR were also included in Figure 1A. One sided paired student t test; *p < 0.05, **p < 0.01, ****p < 0.0001.

### Identification of mTOR-mediated RBP binding to AREs in the IL2 3’UTR

Having established that AREs within cytokine 3’UTRs were required for mTOR-mediated protein production, we next aimed to identify which RBP drives this regulation. As a representative example, focused on the *IL2* 3’UTR, and we performed an RNA aptamer pulldown ^31,36^. We first generated *in vitro* transcribed RNA with streptavidin binding 4×S1 RNA aptamers containing the full-length *IL2* WT 3’UTR, or the *IL2* ARE-mut 3’UTR as bait (Figure 3A). The aptamers were then exposed to cell lysates from *in vitro* generated human CD3^+^ T cells that were activated for 3h with αCD3/αCD28 in the presence or absence of Torin-1. With this approach we could 1) measure RBP binding in the context of the full T cell proteome, 2) define ARE-dependent interactions upon T cell activation, and 3) study the effects of mTOR inhibition on RBP interactions with the target mRNA.

**Figure 3.**
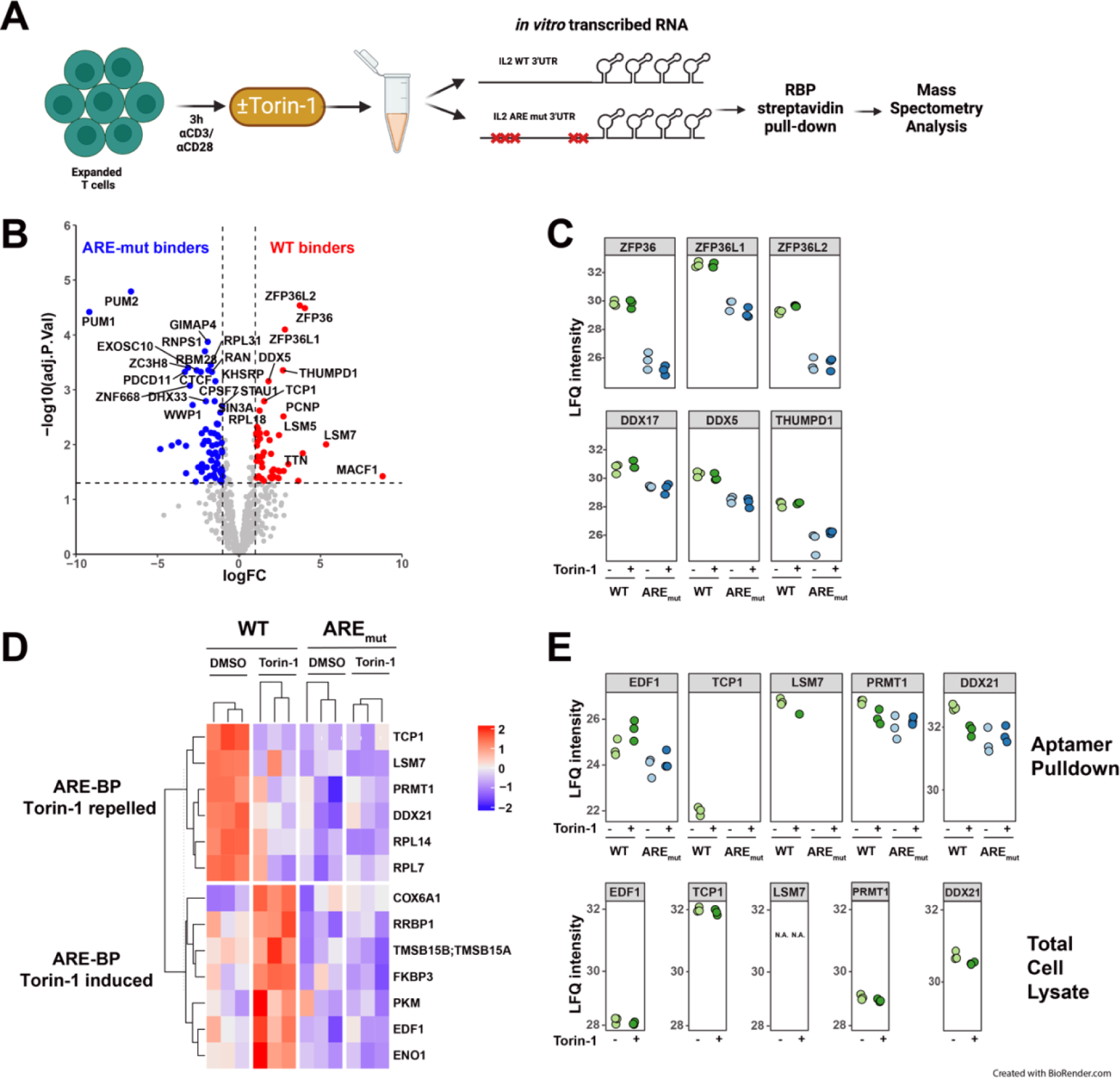
Identification of mTOR-regulated RBPs that interact with the *IL2* 3’UTR. A) Scheme of *IL2* 3’UTR aptamer pulldown to identify mTOR driven ARE binding proteins using *in vitro* transcribed RNA. CD3^+^ Teff cells generated from 40 pooled donors (n=3 donor pools) were activated with aCD3/aCD28 for 3h in the presence of Torin-1 or DMSO. Cell lysates were incubated with *in vitro* transcribed 4xS1m RNA aptamers containing the full length *IL2* WT or *IL2* ARE-mut 3’UTR. Protein binding to 4xS1m RNA aptamers was identified by label-free MS analysis. B) Volcano plot of proteins identified in *IL2* WT or *IL2* ARE-mut in DMSO conditions. Only proteins identified in all three replicates were considered putative ARE binders. Red dots represent proteins that were significantly enriched in the presence of the *IL2* WT 3’UTR; blue dots represent proteins enriched in *IL2* ARE-mut 3’UTR. C) LFQ intensities of top ARE-binding hits from (B) shown in *IL2* WT 3’UTR (green) or *IL2* ARE-mut 3’UTR (blue) in Torin-1 and DMSO conditions. (n=3 donor pools) D) Heatmap of supervised classification displaying proteins that were enriched in *IL2* WT 3’UTR under DMSO conditions (ARE-BP Torin repelled), or proteins enriched in *IL2* WT 3’UTR under Torin-1 conditions, (ARE-BP Torin-1 induced). Correlation between hits and disorder specific theoretical protein profiles had a pearson correlation of >0.75. E) LFQ intensities of identified ARE-BPs that were ARE-BP Torin repelled or ARE-BP Torin-1 induced in (D). Top: Aptamer Pulldown Below: Total Cell Lysate. (n= 3 donor pools). Two-sided t test; FDR = 0.05, S0 = 0.4. na: not applicable. Also see Supplemental Figure 3 and Supplemental Tables 1, 2, 3.

Mass spectrometry analysis of the RNA aptamer pulldown with streptavidin beads detected in total 693 proteins (Supplemental Figure 3A-D & Supplemental table 1). To identify ARE-binding proteins (ARE-BPs), we focused on proteins that showed enriched binding to the *IL2* WT 3’UTR compared to the *IL2* ARE-mut 3’UTR (Figure 3B). As expected, we identified the canonical ARE-BPs ZFP36, ZFP36L1 and ZFP36L2 (Figure 3B), which all depend on AREs for their interaction (Figure 3C). Other previously undescribed ARE interactors included DDX17, DDX5 and THUMPD1 (Figure 3B, C & Supplemental table 2). To identify ARE-BPs that respond to mTOR signaling, we used supervised classification (Supplemental Figure 4E) ^38^. We ordered ARE-BPs with reduced (‘Torin-1 repelled’) or increased (‘Torin-1 induced’) interaction with AREs upon Torin-1 treatment (LFC>1, p<0.05). We found 13 proteins that interacted with AREs in a Torin-1 dependent manner (Figure 3D & Supplemental Figure 3D). Notably, this did not include ZFP36, ZFP36L1, and ZFP36L2, whose interaction to AREs was insensitive to Torin-1 treatment (Figure 3C & Supplemental Figure 3D, E). 6 of the 13 Torin-1 sensitive ARE-BPs (TCP1, LSM7, PRMT1, DDX21, RPL14 and RPL7) displayed reduced interaction, and 7 proteins (COX6A1, RRBP1, TMSB15A/B, FKBP3, PKM, EDF1, ENO1) displayed increased interaction with AREs upon mTOR inhibition (Figure 3D, E top). Of note, none of the observed changes in binding of AREs upon Torin-1 treatment could not be attributed to overall changes in protein expression, which remained unaltered (Figure 3E, bottom). In conclusion, we show here that in addition to the classical (mTOR-insensitive) ARE-BPs - RBPs can depend on mTOR signaling for their interaction to AREs.

### mTOR regulates IL2 cytokine expression through the 3’UTR-binding protein DDX21

We followed up on 5 proteins that we identified as mTOR-sensitive ARE-BPs, and that displayed increased (EDF1) or decreased (TCP1, PRMT1, LSM7, and DDX21) interaction with the *IL2* 3’UTR upon Torin-1 treatment (Figure 3D, E). Upon successful CRISPR-Cas9 gene-editing (Figure 4A), all RBP deficient CD3^+^ T cells – except for EDF1 – had lower cell viability and lower cell count compared to non-targeting CRISPR RNA control, highlighting their relevance for T cell fitness (Supplemental Figure 4A, B). Most strongly affected were LSM7 KO T cells and TCP1 KO T cells, the latter as reported for murine T cells ^39^.

In non-activated ARE-BP deficient T cells, GFP protein expression were comparable to that of control T cells. Also Torin-1 treatment had no noticeable effects on protein expression in resting T cells. Upon T cell activation, however, only EDF1 KO T cells maintained similar GFP expression levels as the control (Figure 4B). Deletion of all other ARE-BP candidates resulted in 30-35% reduction of the GFP gMFI expression compared to control treated T cells, indicating that TCP1, LSM7, PRMT1 and DDX21 regulate protein expression through the *IL2* 3’UTR (Figure 4B). Of note, all RBP-KO cells reached similar induction levels of the activation marker CD69, indicating that not all features of T cell activation were affected (Supplementary Figure 4C).

**Figure 4.**
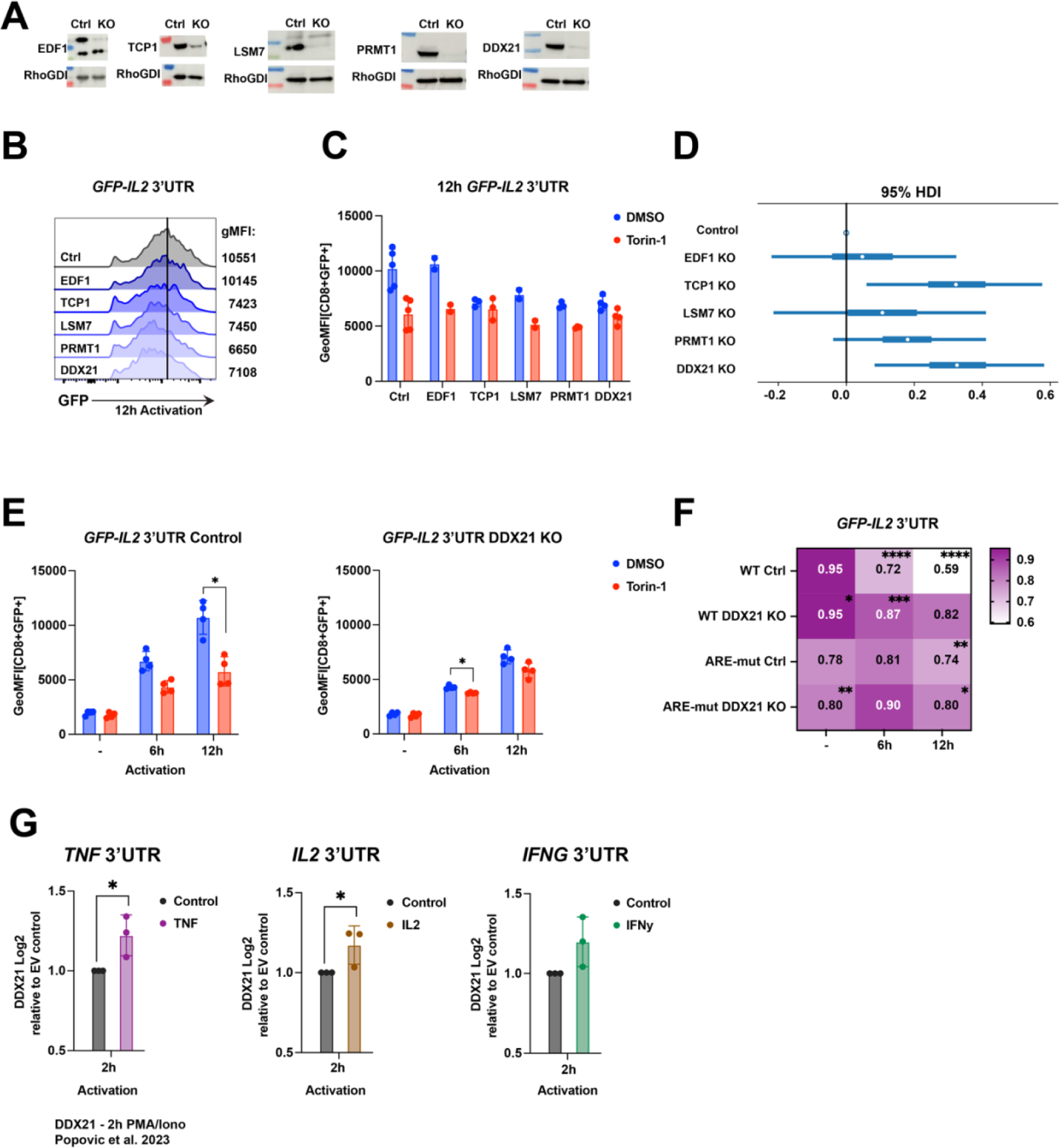
mTOR promotes protein expression through the 3’UTR-binding protein DDX21. A) Immunoblot of CD3^+^ Teff cells 6 days after CRISPR-Cas9 gene editing for indicated RBP, or with non-targeting crRNA (ctrl). B) –D) RBP knockout or non-targeting crRNA (Ctrl) CD3^+^ Teff cells expressing *GFP-IL2* 3’UTR were activated with PMA/Ionomycin. (B) Representative histogram of GFP expression at 12h post activation. geoMFI indicated for each sample at the right. (C) geoMFI of Torin-1 (red) or DMSO (blue) treated cells. (-) indicates non-activated T cells treated with Torin-1 or DMSO for 12h. (n=3-6 donors, mean ± SD). (D) Corresponding Forest plot with Bayesian parameter estimates for the mean DMSO/Torin-1 fraction differences between RBP KO cells calculated with 95% credible intervals. E) -F) CD8^+^ Teff cells expressing *GFP*-*IL2* 3’UTR or *GFP-IL2* ARE-mut 3’UTR and gene-edited for DDX21 knockout or non-targeting crRNA (Ctrl) were activated with PMA/Ionomycin in the presence of Torin-1 or DMSO. (E) geoMFI of GFP plotted for time points. (F) Heatmap scale representative of GFP ratio between DMSO/Torin-1 treated cells. GFP *control* and *GZMB*-3’UTR served as control. (n=2-4 donors, mean ± SD, compiled from 2 independently performed experiments) G) Log2 intensity levels of DDX21 protein binding to the full length 3’UTRs of human *TNF, IL2, IFNG* relative to *Empty Vector (EV)* control in CD3^+^ Teff cells that were activated for 2h with PMA/Ionomycin. (n=3 donor pools). Data extracted from Popovic *et al.* ^31^ E-G: One sided paired student t test; *p < 0.05, **p < 0.01, ***p < 0.001, ****p < 0.0001. Also see Supplemental Figure 4.

We then tested the effect mTOR inhibition on activated ARE-BP KO T cells. EDF1 KO and LSM7 KO T cells maintained their sensitivity to Torin-1 treatment, which was also observed to some extent for PRMT1 KO T cells (Figure 4C). In contrast, TCP1 KO and DDX21 KO T cells almost completely lost the sensitivity to Torin-1 treatment (Figure 4C, D). Because TCP1 deficiency substantially affected cell viability and expansion (Supplemental Figure 4A, B), we focused our studies on DDX21.

Measuring protein expression levels over time revealed that DDX21 KO T cells phenocopied the GFP expression levels of Torin-1 treated control T cells (Figure 4E), indicating that DDX21 is a prominent contributor to the mTOR-mediated, *IL2* 3’UTR-driven protein production. Furthermore, DDX21 deficient cells phenocopied the protein expression levels of *IL2* 3’UTR ARE mutants, and DDX21 KO T cells did not further alter protein expression in the presence of the *IL2* 3’UTR ARE mutant (Figure 4F), providing additional evidence that the prime activity of DDX21 on GFP protein expression is through AREs in the *IL2* 3’UTR.

To study if DDX21 is phosphorylated downstream of mTOR signaling, we mined phospho-proteomics data of activated Jurkat cells ^40^ and mouse T cells ^10^. Both studies identified several species-specific and conserved activation-induced serine/threonine phosphorylation sites in DDX21 (Supplementary Figure 4D). Notably, some phosphorylation sites were lost in T cells from Raptor KO mice (Supplementary Figure 4D), further indicating that phosphorylation of DDX21 is downstream of mTOR signaling.

Finally, to determine whether DDX21 could be involved also in the observed mTOR-mediated regulation of TNF and IFNγ protein production, we examined its binding to the *TNF* 3’UTR and the *IFNG* 3’UTR. Using our previously published aptamer-pulldown data with the three cytokine 3’UTRs from activated T cells^31^ we found that DDX21 displayed significantly increased interaction for the *TNF* 3’UTR and *IL2* 3’UTR when compared to the empty aptamer control, and a trend for increased interaction with the *IFNG* 3’UTR (Figure 4G). Thus, the observed mTOR-mediated interaction of the novel ARE-BP DDX21 to 3’UTRs is observed upon T cell activation for all three measured cytokines.

## DISCUSSION

The interplay between mTOR signaling and cytokine regulation represents a crucial nexus for controlling immune responses. mTOR impacts the availability and activity of translation initiation factors, nutrients and ribosomal proteins ^3,41^. In addition, mTOR signaling can regulate mRNA stability and translation rate, by modulating the binding of LARP to the 5’UTR of its target mRNAs ^20–22,42–44^. Here, we uncovered a previously uncharacterized 3’UTR-mediated mechanism that mTOR employs to promote cytokine production upon T cell activation: mTOR signaling promotes binding of DDX21 to AREs in the cytokine 3’UTRs, which in turn supports the cytokine production. To our knowledge, this is the first report that reveals a link between T cell activation-induced mTOR signaling and ARE dependent 3’UTR-mediated control of protein production.

We here identified mTOR-sensitive and mTOR-insensitive ARE-BPs. The prototypic ARE-BPs such as ZFP36 family members were mTOR-insensitive. ARE-BPs that were mTOR sensitive included DDX21. Intriguingly, the mTOR-mediated interaction of DDX21 with AREs measurable already at 2-3h upon T cell activation promotes protein expression. In contrast, ZFP36 family members dampen cytokine production by promoting RNA degradation ^47^. However, they do so by interacting with AREs at later time points, i.e. from 4h onwards of T cell activation ^31,48,49^. Thus, AREs appear to be prime hubs for fine-tuning the regulation of cytokine production. It is yet to be determined whether the replacement of DDX21 by ZFP36 proteins is passive, dictated by loss of phosphorylation of DDX21, or whether ZFP36 family members have a higher abundance or an affinity for AREs, and thus outcompete DDX21 for binding to AREs. Irrespective of the underlying mechanism, we show here that mTOR mediated regulation of DDX21 binding supports the rapid cytokine production upon T cell activation, a feature that is critical to dampen viral spread.

Not all three cytokine 3’UTRs respond equally strongly to Torin-1 treatment. This different sensitivity could possibly stem from the location of the ARE motifs within the 3’UTR. In the *IFNG* 3’UTR, AREs are located at the 5’end, in the *TNF* 3’UTR at the 3’end, and in the *IL2* 3’UTR at both ends. Provided that 3’UTRs can be shortened upon T cell activation ^24,50,51^, alternative 3’UTR splicing could possibly remove AREs when located at the 3’end. Therefore, TNF may depend less on ARE-mediated regulation and more on other regulatory features, such as stem-loop -mediated regulation ^36^. Furthermore, we recently reported that the relative contribution to protein abundance of sequence features such as CTTT and ARE motifs strongly depends on their location, occurrence and spacing in the mRNA, and this relation holds true for 3’UTRs ^52^. Therefore, it is tempting to speculate that the sequence context of AREs in the three cytokine 3’UTRs (and potentially that of other ARE-containing 3’UTRs) determines the sensitivity to mTOR signaling.

mTOR is considered a key regulator of translation. However, due to low expression levels of the inhibitory protein 4E-BP in human T cells, which results in a stoichiometry between 4E-BP and eIF4E of 3-8:1, it is postulated that only a subset of eIF4E-sensitive transcripts, including proinflammatory effector molecules, are responsive to 4E-BP in T cells and consequently to mTOR regulation ^10,12,13,53,54^. We found here that cytokine 3’UTRs can substantially contribute to the mTOR-mediated protein production upon T cell activation. This was not the case for *GZMB* 3’UTR, possibly because the human *GZMB* 3’UTR contains only one ARE motif.

In conclusion, we present a novel mechanism that mTOR employs to control protein production. Our results reveal the effects of mTOR signaling on determining RBP binding to AREs within the 3’UTR and, consequently, on mRNA processing and translation. The ARE-mediated effects of mTOR signaling thus contribute to the regulation of pro-inflammatory cytokine production, which is crucial for coordinating and regulating T cell responses in health and disease.

## RESOURCE AVAILABILITY

Lead contact Further information and requests for resources and reagents should be directed to and will be fulfilled by the lead contact, Monika C. Wolkers.

## MATERIALS AVAILABILITY

This study did not generate new unique reagents.

## DATA CODE AND AVAILABILITY

- The mass spectrometry proteomics data of the RNA Aptamer Pulldown have been deposited to the ProteomeXchange Consortium via the PRIDE^55^ partner repository with the dataset identifier PXD056415
- This paper does not report original custom code. All codes used in this paper are available from the lead contact upon request.
- Any additional information required to reanalyze the data reported in this paper is available from the lead contact upon request.

## ACKNOWLEDGMENTS

We thank G. Stoecklin (University of Heidelberg) for the 4xS1m aptamer construct; L. Wardak and A. Guislain for technical help; V. Lattanzio, N. Zandhuis, K. Bresser, for critical reading of the manuscript. This research was supported by European Research Council consolidator award PRINTERS 817533, and Oncode Institute, all to M.C.W.

## AUTHOR CONTRIBUTIONS

Conceptualization, A.P.J and M.C.W.; Methodology, A.P.J., B.P., and M.C.W.; Investigation, A.P.J., B.P., J.Z., and F.P.J.v.A.; Formal Analysis, A.P.J., A.J.H., and K.R.; Validation, A.P.J., J.Z., and A.B.; Writing-Original Draft, A.P.J and M.C.W.; Writing – Review & Editing, A.P.J., B.P., A.J.H., F.P.J.v.A., and M.C.W.; Supervision, M.C.W.; Funding Acquisition, M.C.W.

## METHODS

### Cell culture

Peripheral blood mononuclear cells (PBMCs) from anonymized healthy donors were used in accordance with the Declaration of Helsinki (Seventh Revision, 2013) after written informed consent (Sanquin). PBMCs were isolated through Lymphoprep density gradient separation (Stemcell Technologies). Cells were used after cryopreservation. T cells were activated for 48h as described previously ^56^. Briefly, 24-well plates were pre-coated overnight at 4°C with 4µg/mL rat a-mouse IgG2a (MW1483, Sanquin) in phosphate-buffered saline (PBS). Plates were washed with PBS and coated for >3 h with 1 µg/mL αCD3 (HIT3a, Biolegend) at 37°C. 1.3×10^6^ PBMCs/well were seeded with 1 µg/mL soluble αCD28 (CD28.2, Biolegend) in 1 mL Iscove’s Modified Dubecco’s Medium (IMDM) supplemented with 10% fetal bovine serum (FBS), 100 U/mL penicillin, 100 µg/mL streptomycin, and 2 mM L-glutamine. After 48h of activation at 37°C, 5% CO2, cells were harvested and cultured in standing T25/T75 tissue culture flasks (Thermo Scientific) at a density of 0.8×10^6^/mL in IMDM supplemented with 10% FBS and 100 IU/mL recombinant human rhIL2 (Protech) and 10 ng/mL rhIL-15 (Protech). Medium was refreshed every 2-3 days. Upon nucleofection, T cells were cultured in T cell mixed media (Miltenyi) supplemented with 5% heat-inactivated human serum, 5% FBS, 100 U/mL Penicillin, 100 mg/mL streptomycin, 2 mM L-glutamine, 100 IU/mL rhIL-2 (Peprotech), and 10 ng/mL rhIL-15 (Peprotech). Human fibrosarcoma FLYRD18 cells (FLYRD18) were cultured in IMDM supplemented with 10% FBS, 1% L-glutamine, 1% Penicillin-Streptomycin antibiotic solution at 37°C, 5% CO2 and split every 2 days at 1:10.

### Generation of GFP reporter constructs

GFP constructs with full length cytokine 3’UTRs were reported before ^31^. Cytokine 3’UTR fragments and full length 3’UTRs containing CTTT--> CGTT or AUUUA--> AUGUA point mutants containing BamHI and NotI restriction sites were obtained as minigenes from IDT and inserted into pRS_PGK_GFP according to manufacturer’s protocol (T4; NEB). Plasmid DNA was amplified in DH5alpha super-competent bacteria (NEB) and isolated with NucleoSpin Plasmid Transfection-grade Mini-Prep isolation kit (Machery-Nagel). 3’UTR sequences were confirmed with Sanger sequencing.

### Virus production and retroviral transduction

1×10^5^ FLYRD18 cells per 6-well culture plates were pre-seeded and cultured overnight at 37°C. Transfection with GFP reporter constructs was carried out with GeneJammer (Agilent) according to the manufacturer’s protocol and incubated overnight at 32°C. T cells activated for 48h were retrovirally transduced with Retronectin (Takara) as previously described ^31^. Briefly, non-tissue cultured treated 24-well plates were pre-coated overnight with 50 µg/mL Retronectin (Takara) and washed once with PBS. 800 µL viral supernatant/well was added and spun at 4°C at 2820g. 1×10^6^ T cells/well were added, plates spun for 5 min at 180g and incubated overnight at 37°C. Supernatant was replaced with fresh culture medium. After 48h, T cells were harvested and cultured in T25/T75 flasks for 6–8 days as described above.

### T cell activation and flow cytometric measurement of protein expression

T cells were activated in culture medium in 96-wells plates with 10 ng/mL Phorbol Myristate Acetate (PMA) and 1μM Ionomycin, or with αCD3 (Pelicluster; Sanquin) and αCD28 (Biolegend), 1µg/mL each. To block mTOR, 250nM Torin-1, or 10nM Rapamycin or dissolvent DMSO (0.05% for Torin-1, 0.01% for Rapamycin) was added during stimulation. Non-stimulated T cells treated with mTOR inhibitors and/or DMSO control for 12h were included as controls.

To analyze GFP protein expression, T cells were washed with FACS buffer (PBS, containing 1% FBS and 2 mM EDTA) and labeled for 20 min at 4°C with indicated antibodies (see Appendix A). Dead cells were excluded with Near-IR (Life Technologies). For intracellular cytokine staining, cells were cultured with 1 µg/mL brefeldin A for indicated time points, fixed, and permeabilized with Cytofix/Cytoperm kit according to the manufacturer’s protocol (BD Biosciences) prior to acquisition using FACSymphony. Data were analyzed with FlowJo (BD Biosciences, version 10).

### FluoroSpot Cytokine Assay

Fluorospot cytokine assay was performed at day 4 after T cell activation according to the manufacturer’s instructions (Mabtech). Briefly, 1×10^4^ CD3^+^ T cells/well were plated in pre-coated IFNγ/IL2/TNF Fluorospot plates (Mabtech) and cultured overnight in medium alone (negative control) or were activated with αCD3/αCD28 (supplied by kit), in the presence of 250nM Torin-1 or DSMO control 12h. Fluorospot plates were developed and evaluated using the ImmunoSpot S6 Ultra-V (CTL) with the accessory software.

### Quantitative PCR analysis

Total RNA was extracted using Quick-RNA MiniPrep plus kip (Zymo). T cells were activated in triplicate for indicated 3h with αCD3/αCD28. For mRNA half-life measurements, cells were treated for an additional 1h or 2h with 5 µg/mL actinomycin D (ActD) (Sigma-Aldrich). cDNA was synthesized with SuperScript III (Invitrogen), RT-PCR was performed using SYBR Green on a StepOne Plus (Applied Biosystems). Ct values were normalized to 18S levels.

### Generation of in vitro transcribed S1m aptamers containing the *IL2* 3’UTR

Full-length 3’UTRs were ordered as gBlocks from IDT and cloned into BamHI and NotI sites of pRETRO-SUPER-GFP downstream of GFP ^31^. 4xS1m RNA aptamers containing the full-length human WT *IL2* 3’UTR or ARE-mut *IL2 3’*UTR were cloned into the pSP73-flip-4xS1m vector, linearized with NdeI, and *in vitro* transcribed with AmpliScribe T7 flash transcription kit (Epicentre) as previously described ^18,31^. RNA quality and quantity was determined by RNAnano Chip assay (Agilent).

### 4xS1m RNA aptamer-protein pull-down

CD3^+^ T cells were activated for 48h with αCD3/αCD28 and rested for 5 days in the presence of 100 IU/mL rhIL2 as described above. T cells were left untreated or reactivated for 3h with αCD3/αCD28 in the presence of Torin-1 or DMSO. Cells were pelleted and washed twice with ice-cold PBS. Cell pellet was snap frozen in liquid nitrogen, homogenized using 5-mm steel beads and a tissue lyser (Qiagen TissueLyser II) 6x at 25 Hz for 15s. The homogenate was solubilized and precleared with Avidin agarose beads (Thermo Scientific) for 30 min at 4°C and with Streptavidin Sepharose High Performance beads (GE Healthcare) for 2h at 4°C. Cell lysates were incubated with RNA-aptamer-coupled beads for 3.5h at 4°C under rotation in the presence of 60 U RNasin (Ambion). For each pull-down, 30 µg of in vitro transcribed RNA, coupled to Streptavidin Sepharose beads, and 5–10mg cell lysate protein was used. RNA-bound proteins were eluted by adding 1 µg RNaseA (Thermo Scientific) in 100mL 100mM Tris-HCl, pH 7.5 (Gibco-Invitrogen). Proteins were reduced, alkylated, and digested into peptides using trypsin (Promega). Peptides were desalted and concentrated using Empore-C18 StageTips and eluted with 0.5% (v/v) acetic acid, 80% (v/v) acetonitrile. Sample volume was reduced by SpeedVac and supplemented with 2% acetonitrile and 0.1% TFA.

### Mass spectrometry data acquisition

Tryptic peptides were separated by nanoscale C18 reverse phase chromatography coupled online to an Orbitrap Fusion Lumos Tribrid mass spectrometer (Thermo Scientific) via a nanoelectrospray ion source (Nanospray Flex Ion Source, Thermo Scientific). Peptides were loaded on a 20cm 75–360µm inner-outer diameter fused silica emitter (New Objective) packed in-house with ReproSil-Pur C18-AQ, 1.9μm resin (Dr Maisch GmbH). The column was installed on a Dionex Ultimate3000 RSLC nanoSystem (Thermo Scientific) using a MicroTee union formatted for 360μm outer diameter columns (IDEX) and a liquid junction. The spray voltage was set to 2.15kV. Buffer A was composed of 0.1% formic acid and buffer B of 0.1% formic acid, 80% acetonitrile. Peptides were loaded for 17 min at 300nl/min at 5% buffer B, equilibrated for 5 min at 5% buffer B (17-22 min) and eluted by increasing buffer B from 5-28% (22-80 min) and 28-40% (80-85 min), followed by a 5 min wash to 95 % and a 5 min regeneration to 5%. Survey scans of peptide precursors from 375 to 1500 m/z were performed at 120K resolution (at 200 m/z) with a 4×10^5^ ion count target. Tandem mass spectrometry was performed by isolation with the quadrupole with isolation window 0.7, HCD fragmentation with normalized collision energy of 30, and rapid scan mass spectrometry analysis in the ion trap. The MS2 ion count target was set to 3×10^4^, and the max injection time was 20ms. Only those precursors with charge state 2–7 were sampled for MS2. The dynamic exclusion duration was set to 20s with a 10ppm tolerance around the selected precursor and its isotopes. Monoisotopic precursor selection was turned on. The instrument was run in top speed mode with 1s cycles. All data were acquired with Xcalibur SII software.

### Mass spectrometry analysis

Raw mass spectrometry files were processed with the MaxQuant computational platform, version 1.6.2.10. Proteins and peptides were identified using the Andromeda search engine by querying the reviewed human Uniprot database (downloaded 09-22-2023 20426 entries). Standard settings with the additional options match between runs, and unique peptides for quantification were selected. The generated ‘proteingroups.txt’ data were imported in R4.30/Rstudio 2024.04.1. ‘reverse’, ‘potential contaminants’ and ‘only identified by site’ peptides were filtered out and label free quantification values were log2 transformed.

Missing values were imputed by a normal distribution (width=0.3, shift = 1.8), assuming these proteins were close to the detection limit. Statistical analyses were performed using moderated t-tests in the LIMMA package. A Benjamini-Hochberg adjusted P value <0.05 and absolute log2 fold change >1 was considered statistically significant and relevant.

For supervised classification, we generated theoretical profiles in which conditions of interest intensities were set to an arbitrary high and low in all other samples ^38^. These profiles were correlated with all statistically significant proteins. Correlation between data and disorder specific theoretical protein profiles was performed using a Pearson correlation coefficient of 0.5 and Benjamini-Hochberg adjusted P value <0.05 was used as threshold. The R package ggplot2 was used for graphical representations.

### Genetic modification of T cells with Cas9 RNPs

crRNAs were designed in Benchling (https://benchling.com; Appendix B). Cas9 RNP production and T cell nucleofection was performed as previously described ^35^. Briefly, Alt-R crRNA and tracrRNA were reconstituted to 100 µM in Nuclease Free Duplex buffer (all IDT). As a negative control, nontargeting negative control crRNA #1 was used (IDT). Oligos were mixed at equimolar ratios (i.e. 4.5 mL total crRNA + 4.5 mL tracrRNA) in nuclease-free PCR tubes and denatured by heating at 95°C for 5 min in a thermocycler. Nucleic acids were cooled down to room temperature prior to mixing them with 30 µg Alt-R™ S.p. Cas9 Nuclease V3 (IDT) to produce Cas9 ribonuclear proteins (RNPs). Mixture was incubated at room temperature for at least 10 min prior to nucleofection. For nucleofection, human CD3^+^ T cells were activated for 48 h with αCD3/αCD28 and washed twice with PBS. Cells were resuspended in P2 buffer (Lonza). Cells were electroporated in 16-well strips in a 4D Nucleofector X unit (Lonza) with program EH100 for human T cells &P2 buffer. Knockout efficiency was determined on day 5 days after electroporation by Western blot.

### Immunoblotting

Cell lysates (1×10^6^ cells per sample) were prepared by standard procedures using RIPA lysis buffer (Thermo) supplemented with protease and phosphatase inhibitors (Thermo). Lysates were run on 4–12% SDS-PAGE (Thermo). SDS-PAGE gels were directly transferred onto nitrocellulose membranes (iBlot2, Thermo). Subsequently, membranes were blocked with 5% BSA TBST solution (Fraction V, Sigma). Membranes were stained with α-EDF1 (Proteintech), α-TCP1 (91A, Thermo Fisher Scientific) α-LSM7 (Proteintech), α-PRMT1 (A33, Cell Signalling Technology), α-DDX21 (Proteintech) (or α-RhoGDI (MAB9959, Abnova), followed by α-Rabbit (4050-05, Southern Biotech) or α-Mouse (1031-05, Southern Biotech) HRP-conjugated secondary antibodies.

## Quantification and statistical analysis

Results are shown as mean ± SD. Statistical analysis was performed with GraphPad Prism 10 with Wilcoxon two-tailed ratio paired or unpaired Student’s t test when comparing two groups, or with one-way ANOVA test with Dunnett correction when comparing more than two groups. p values < 0.05 were considered statistically significant.

## Appendix: Key resources

### Appendix A

| Antibodies | Sources |
| --- | --- |
| Rat anti-mouse IgG2a (clone MW1483) | Sanquin |
| Mouse anti-CD28 (clone CD28.2) | Biolegend Cat# 302902 |
| Mouse anti-CD3 (clone HIT3a) | Biolegend Cat# 300302 |
| Pelicluster CD3 | Sanquin Cat#564364 |
| BUV737 mouse anti-human CD4 (clone SK3) | BD Bioscience Cat# 564305 |
| BUV805 mouse anti-human CD8 (clone SK1) | BD Bioscience Cat# 612889 |
| BUV395 mouse Anti-human CD69 (clone FN50) | BD Bioscience |
| Brilliant Violet 785™ anti-human TNF (clone MAb11) | Biolegend Cat# 502948 |
| eFluor 450 mouse anti-human IFN gamma (clone 4S.B3) | eBiosciences Cat# 48-7319-42 |
| eFluor 450 mouse anti-human CD3 (clone OKT3) | eBioscience Cat#48-0037-42 |
| APC mouse anti-human IL2 (clone MQ1-17H12) | Biolegend Cat# 500314 |
| LIVE/DEAD™ Fixable Near-IR Dead Cell Stain Kit, for 633 or 635 nm excitation | Invitrogen Cat# L34976 |
| CellTrace Violet© | Thermo Fisher Scientific |
| Alexa Fluor® 647 mouse anti-human CD137 (clone 4B4-1) | Biolegend Cat# 309824 |
| TCP1 rat anti-human monoclonal antibody (91A) | Thermo Fisher Scientific Cat# MA3-026 |
| DDX21 rabbit anti-human Polyclonal antibody | Proteintech cat# 10528-1-AP |
| EDF1 rabbit anti-human Polyclonal | Proteintech cat# 12419-1-AP |
| PRMT1 rabbit anti-human antibody (A33) | Cell Signaling Technology Cat# 2449S |
| LSM7 rabbit anti-human Polyclonal antibody | Proteintech Cat# 18941-1-AP |
| Mouse anti-RhoGDI | Abnova Cat# 89-113-917 |
| **Bacterial and viral strains** |  |
| E. coli DH5alpha super competent bacteria | Invitrogen Cat#1 8265017 |
| **Biological samples** |  |
| Human PBMC | Sanquin |
| Human: FLYRD18 cells | ECACC |
| **Chemicals, peptides, and recombinant proteins** |  |
| Trypsin Gold, Mass Spectrometry Grade | Promega Cat# V5280 |
| GolgiPlug Protein Transport Inhibitor (containing Brefeldin A) | BD Biosciences Cat# BD 555029 |
| Recombinant human IL-15 | Peprotech Cat# 200-15 |
| Recombinant human IL2 | Peprotech Cat# 200-02 |
| GeneJammer | Agilent Cat# 204130 |
| RetroNectin | Takara Bio Cat# T100B |
| Phorbol 12-myristate 13-acetate (PMA) | Sigma-Aldrich Cat# P8139 |
| Ionomycin calcium salt | Sigma-Aldrich Cat# 13909 |
| RNase A | Thermo Scientific Cat# EN0531 |
| Precision Counting Beads | Biolegend Cat#424902 |
| GelRed Nucleic Acid Gel Stain | Biotium Cat# 41003 |
| Actinomycin D | Sigma-Aldrich Cat# A9415 |
| Alt-R™ S.p. Cas9 Nuclease V3 | IDT Cat# 10000735 |
| Power SYBRTM Green PCR Master Mix | Applied Biosystems Cat# 4367660 |
| RNase A, DNase and protease-free (10 mg/mL) | Thermo Scientific Cat# EN0531 |
| RNaseOUT™ Recombinant Ribonuclease Inhibitor | Invitrogen Cat# 10777019 |
| Ribonucleoside Vanadyl Complex | New England Biolabs Cat# S1402S |
| Halt™ Protease and Phosphatase Inhibitor Single-Use Cocktail, EDTA-Free (100X) | Thermo Scientific Cat# 78443 |
| Digitonin, High Purity | Calbiochem Cat# 300410 |
| Lympholyte-M | Cederlane Cat# CL5035 |
| Dynabeads™ Protein A for Immunoprecipitation | Invitrogen Cat# 10008D |
| Dynabeads™ Protein G for Immunoprecipitation | Invitrogen Cat# 10003D |
| Streptavidin Sepharose® High Performance | GE Healthcare Cat# GE17-5113-01 |
| **Critical commercial assays** |  |
| NucleoSpin Plasmid Transfection grade, Mini kit for ultrapure plasmid DNA | Machery-Nagel |
| Fixation/Permeabilization Solution Kit with BD GolgiPlug™ | BD Biosciences Cat# 555028 |
| P2 Primary Cell 4D-Nucleofector™ X Kit S | Lonza Cat# V4XP-2032 |
| SuperScript™ III First-Strand Synthesis System | Invitrogen Cat# 18080051 |
| AmpliScribe™ T7-Flash™ Transcription Kit | Epicentre Cat# ASF3507 |
| Quick-RNA MiniPrep plus kip | Zymo Cat# R1058 |
| FluoroSpot Plus: Human IFNγ/IL2/TNF | Mabtech # FSP-010209-2 |
| **Equipment** |  |
| VerityPro Thermal Cycler | Thermo Scientific |
| Mupid One Electrophoresis System | Nippon Genetics Europe |
| NGS platform | Illumina |
| NanoDrop 2000 | Thermo Scientific |
| BD LSR Fortessa Cell Analyzer | BD Biosciences |
| FACS Symphony A5 Cell Analyzer | BD Biosciences |
| Mabtech IRIS 2 | Mabtech |
| Amersham ImageQuant 800 | Cytiva |
| Casy Cell counter | Roche Innovatis |
| **Software and algorithms** |  |
| FlowJo version 10.8.1 | BD Biosciences |
| GraphPad Prism version 9.1.1 | GraphPad Software |

### Appendix B

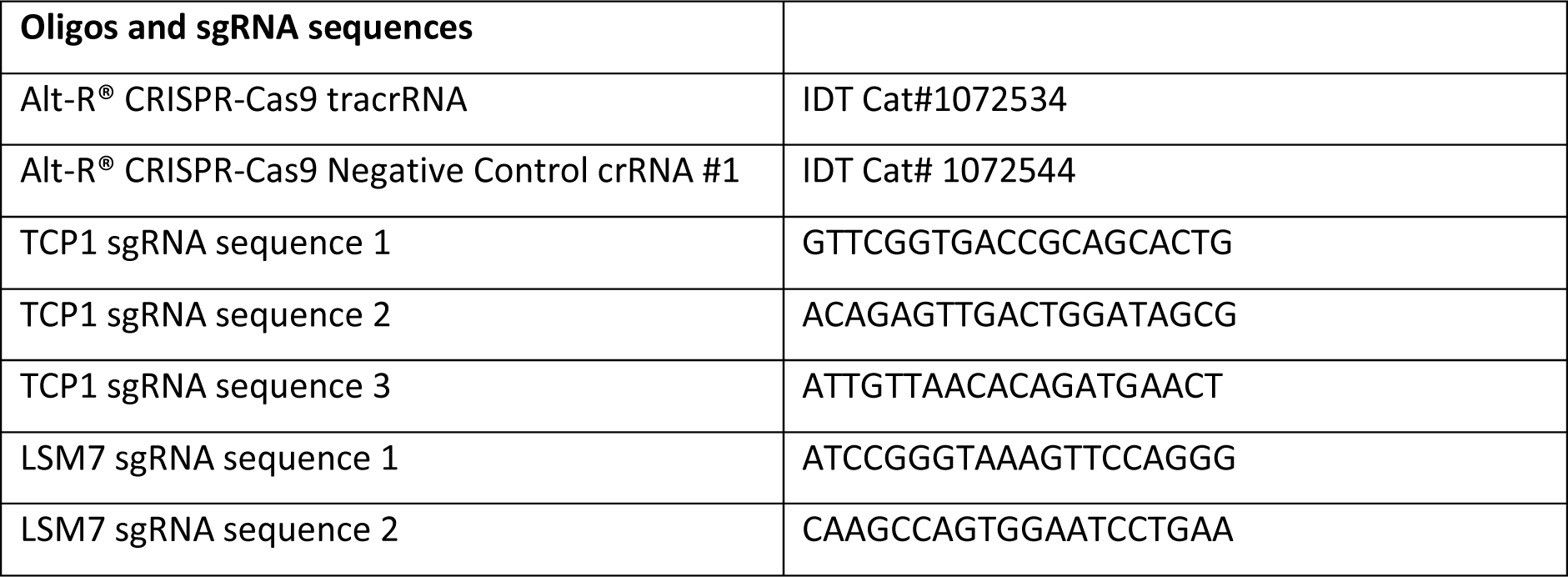
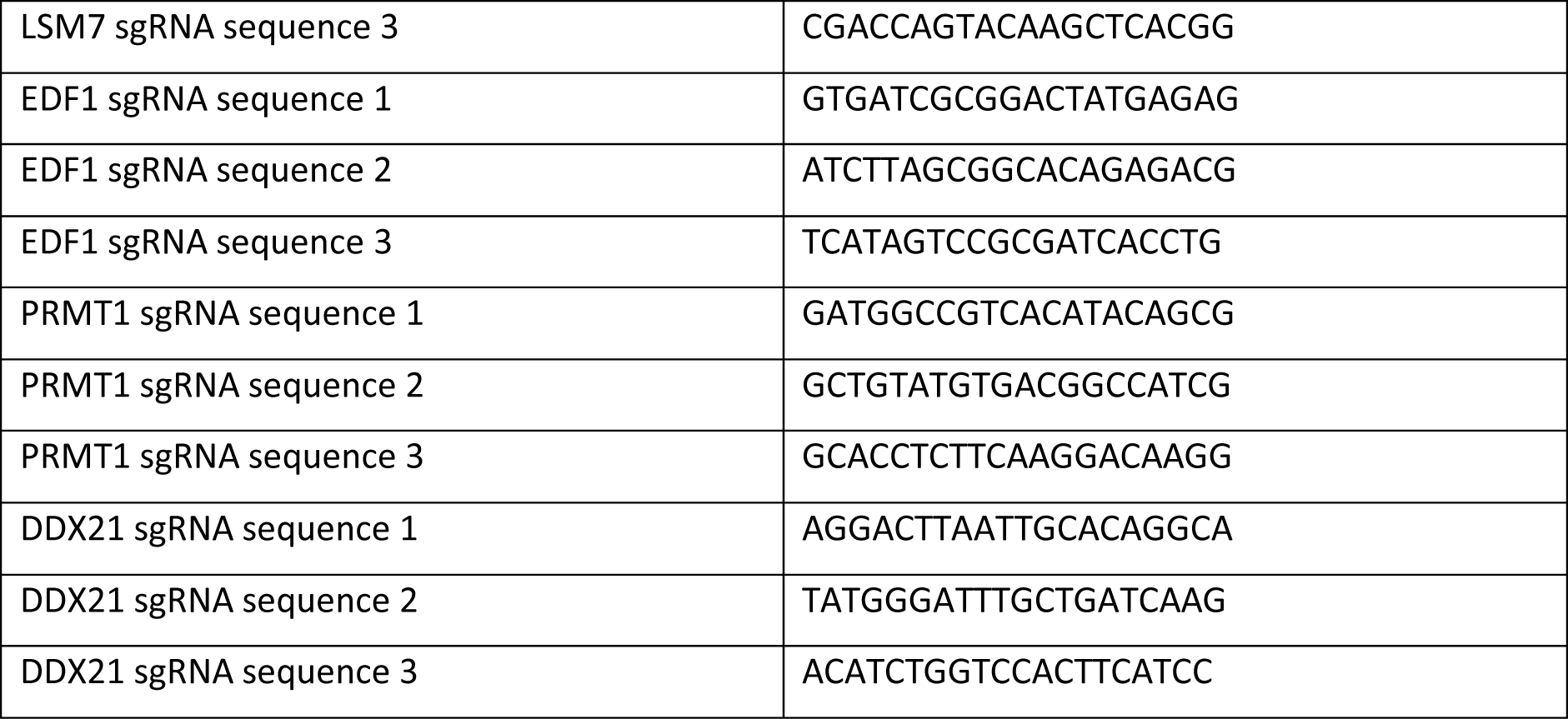

## EXCEL TABLES TITLES

**Supplementary Table 1. *IL2* Aptamer Pulldown results**

**Supplementary Table 2. ARE-binders**

**Supplementary Table 3. Shape analysis top hits**

## References

1. Salerno, F., Turner, M., and Wolkers, M.C. (2020). Dynamic post-transcriptional events governing CD8+ T cell homeostasis and effector function. Trends Immunol. 41, 240–254. 10.1016/j.it.2020.01.001.

2. Nicolet, B.P., Zandhuis, N.D., Lattanzio, V.M., and Wolkers, M.C. (2021). Sequence determinants as key regulators in gene expression of T cells. 1–20. 10.1111/imr.13021.

3. Powell, J.D., and Delgoffe, G.M. (2010). The Mammalian Target of Rapamycin: Linking T Cell Differentiation, Function, and Metabolism. Immunity 33, 301–311. 10.1016/j.immuni.2010.09.002.

4. Chi, H. (2012). Regulation and function of mTOR signalling in T cell fate decisions. Nat. Rev. Immunol. 12, 325–338. 10.1038/nri3198.

5. Pollizzi, K.N., Patel, C.H., Sun, I.H., Oh, M.H., Waickman, A.T., Wen, J., Delgoffe, G.M., and Powell, J.D. (2015). mTORC1 and mTORC2 selectively regulate CD8+ T cell differentiation. J. Clin. Invest. 125, 2090–2108. 10.1172/JCI77746.

6. Hukelmann, J.L., Anderson, K.E., Sinclair, L.V., Grzes, K.M., Murillo, A.B., Hawkins, P.T., Stephens, L.R., Lamond, A.I., and Cantrell, D.A. (2016). The cytotoxic T cell proteome and its shaping by the kinase mTOR. Nat. Immunol. 17, 104–112. 10.1038/ni.3314.

7. Jurgens, A.P., Popović, B., and Wolkers, M.C. (2021). T cells at work: How post-transcriptional mechanisms control T cell homeostasis and activation. Eur. J. Immunol. 51, 2178–2187. 10.1002/eji.202049055.

8. Villa, E., Sahu, U., O’Hara, B.P., Ali, E.S., Helmin, K.A., Asara, J.M., Gao, P., Singer, B.D., and Ben-Sahra, I. (2021). mTORC1 stimulates cell growth through SAM synthesis and m6A mRNA-dependent control of protein synthesis. Mol. Cell 81, 2076–2093.e9. 10.1016/j.molcel.2021.03.009.

9. Schneider, C., Erhard, F., Binotti, B., Buchberger, A., Vogel, J., and Fischer, U. (2022). An unusual mode of baseline translation adjusts cellular protein synthesis capacity to metabolic needs. Cell Rep. 41, 111467. 10.1016/j.celrep.2022.111467.

10. Tan, H., Yang, K., Li, Y., Shaw, T.I., Wang, Y., Blanco, D.B., Wang, X., Cho, J.H., Wang, H., Rankin, S., et al. (2017). Integrative proteomics and phosphoproteomics profiling reveals dynamic signaling networks and bioenergetics pathways underlying T cell activation. Immunity 46, 488–503. 10.1016/j.immuni.2017.02.010.

11. Marchingo, J.M., Sinclair, L.V., Howden, A.J., and Cantrell, D.A. (2020). Quantitative analysis of how myc controls t cell proteomes and metabolic pathways during t cell activation. eLife 9, 1–23. 10.7554/eLife.53725.

12. Wolf, T., Jin, W., Zoppi, G., Vogel, I.A., Akhmedov, M., Bleck, C.K.E., Beltraminelli, T., Rieckmann, J.C., Ramirez, N.J., Benevento, M., et al. (2020). Dynamics in protein translation sustaining T cell preparedness. Nat. Immunol. 21, 927–937. 10.1038/s41590-020-0714-5.

13. Marchingo, J.M., and Cantrell, D.A. (2022). Protein synthesis, degradation, and energy metabolism in T cell immunity. Cell. Mol. Immunol. 10.1038/s41423-021-00792-8.

14. Carballo, E., Lai, W.S., and Blackshear, P.J. (1998). Feedback inhibition of macrophage tumor necrosis factor-alpha production by tristetraprolin. Science 281, 1001–1005. 10.1126/science.281.5379.1001.

15. Kontoyiannis, D., Pasparakis, M., Pizarro, T.T., Cominelli, F., and Kollias, G. (1999). Impaired On/Off Regulation of TNF Biosynthesis in Mice Lacking TNF AU-Rich Elements: Implications for Joint and Gut-Associated Immunopathologies. Immunity 10, 287–398.

16. Ogilvie, R.L., SternJohn, J.R., Rattenbacher, B., Vlasova, I.A., Williams, D.A., Hau, H.H., Blackshear, P.J., and Bohjanen, P.R. (2009). Tristetraprolin Mediates Interferon-γ mRNA Decay. J. Biol. Chem. 284, 11216–11223. 10.1074/jbc.M901229200.

17. Hodge, D.L., Berthet, C., Coppola, V., Kastenmüller, W., Buschman, M.D., Schaughency, P.M., Shirota, H., Scarzello, A.J., Subleski, J.J., Anver, M.R., et al. (2014). IFN-gamma AU-rich element removal promotes chronic IFN-gamma expression and autoimmunity in mice. J. Autoimmun. 53, 33–45. 10.1016/j.jaut.2014.02.003.

18. Salerno, F., Engels, S., van den Biggelaar, M., van Alphen, F.P.J., Guislain, A., Zhao, W., Hodge, D.L., Bell, S.E., Medema, J.P., von Lindern, M., et al. (2018). Translational repression of pre-formed cytokine-encoding mRNA prevents chronic activation of memory T cells. Nat. Immunol. 19, 828– 837. 10.1038/s41590-018-0155-6.

19. Mayr, C. (2019). What are 3′ utrs doing? Cold Spring Harb. Perspect. Biol. 11, 1–16. 10.1101/cshperspect.a034728.

20. Thoreen, C.C., Chantranupong, L., Keys, H.R., Wang, T., Gray, N.S., and Sabatini, D.M. (2012). A unifying model for mTORC1-mediated regulation of mRNA translation. Nature 485, 109–113. 10.1038/nature11083.

21. Hong, S., Freeberg, M.A., Han, T., Kamath, A., Yao, Y., Fukuda, T., Suzuki, T., Kim, J.K., and Inoki, K. (2017). LARP1 functions as a molecular switch for mTORC1-mediated translation of an essential class of mRNAs. eLife 6, 1–24. 10.7554/eLife.25237.

22. Philippe, L., van den Elzen, A.M.G., Watson, M.J., and Thoreen, C.C. (2020). Global analysis of LARP1 translation targets reveals tunable and dynamic features of 5′ TOP motifs. Proc. Natl. Acad. Sci. U. S. A. 117, 5319–5328. 10.1073/pnas.1912864117.

23. Berman, A.J., Thoreen, C.C., Dedeic, Z., Chettle, J., Roux, P.P., and Blagden, S.P. (2021). Controversies around the function of LARP1. RNA Biol. 18, 207–217. 10.1080/15476286.2020.1733787.

24. Chang, J.-W., Zhang, W., Yeh, H.-S., De Jong, E.P., Jun, S., Kim, K.-H., Bae, S.S., Beckman, K., Hwang, T.H., Kim, K.-S., et al. (2015). mRNA 3′-UTR shortening is a molecular signature of mTORC1 activation. Nat. Commun. 6, 7218. 10.1038/ncomms8218.

25. Masopust, D., Vezys, V., Marzo, A.L., and Lefrançois, L. (2001). Preferential Localization of Effector Memory Cells in Nonlymphoid Tissue. Science 291, 2413–2417. 10.1126/science.1058867.

26. Han, Q., Bagheri, N., Bradshaw, E.M., Hafler, D.A., Lauffenburger, D.A., and Love, J.C. (2012). Polyfunctional responses by human T cells result from sequential release of cytokines. Proc. Natl. Acad. Sci. U. S. A. 10.1073/pnas.1117194109.

27. Nicolet, B.P., Guislain, A., and Wolkers, M.C. (2017). Combined Single-Cell Measurement of Cytokine mRNA and Protein Identifies T Cells with Persistent Effector Function. J. Immunol. 198, 962–970. 10.4049/jimmunol.1601531.

28. Thoreen, C.C., Kang, S.A., Chang, J.W., Liu, Q., Zhang, J., Gao, Y., Reichling, L.J., Sim, T., Sabatini, D.M., and Gray, N.S. (2009). An ATP-competitive Mammalian Target of Rapamycin Inhibitor Reveals Rapamycin-resistant Functions of mTORC1. J. Biol. Chem. 284, 8023–8032. 10.1074/jbc.M900301200.

29. Mukhopadhyay, S., Frias, M.A., Chatterjee, A., Yellen, P., and Foster, D.A. (2016). The Enigma of Rapamycin Dosage. Mol. Cancer Ther. 15, 347–353. 10.1158/1535-7163.MCT-15-0720.

30. Zhao, W., and Erle, D.J. (2018). Widespread Effects of Chemokine 3′ Untranslated Regions on mRNA Degradation and Protein Production in Human Cells. J. Immunol. 201, 1053–1061. 10.4049/jimmunol.1800114.

31. Popović, B., Nicolet, B.P., Guislain, A., Engels, S., Jurgens, A.P., Paravinja, N., Freen-van Heeren, J.J., Van Alphen, F.P.J., Van Den Biggelaar, M., Salerno, F., et al. (2023). Time-dependent regulation of cytokine production by RNA binding proteins defines T cell effector function. Cell Rep. 42, 112419. 10.1016/j.celrep.2023.112419.

32. Tcherkezian, J., Cargnello, M., Romeo, Y., Huttlin, E.L., Lavoie, G., Gygi, S.P., and Roux, P.P. (2014). Proteomic analysis of cap-dependent translation identifies LARP1 as a key regulator of 5′TOP mRNA translation. Genes Dev. 28, 357–371. 10.1101/gad.231407.113.

33. Caput, D., Beutler, B., Hartog, K., Thayer, R., Brown-Shimer, S., and Cerami, A. (1986). Identification of a common nucleotide sequence in the 3’-untranslated region of mRNA molecules specifying inflammatory mediators. Proc. Natl. Acad. Sci. 83, 1670–1674. 10.1073/pnas.83.6.1670.

34. Barreau, C. (2005). AU-rich elements and associated factors: are there unifying principles? Nucleic Acids Res. 33, 7138–7150. 10.1093/nar/gki1012.

35. Freen-van Heeren, J.J., Popović, B., Guislain, A., and Wolkers, M.C. (2020). Human T cells employ conserved AU-rich elements to fine-tune IFN-γ production. Eur. J. Immunol., 1–10. 10.1002/eji.201948458.

36. Leppek, K., Schott, J., Reitter, S., Poetz, F., Hammond, M.C., and Stoecklin, G. (2013). Roquin promotes constitutive mrna decay via a conserved class of stem-loop recognition motifs. Cell 153, 869–881. 10.1016/j.cell.2013.04.016.

37. Leppek, K., and Stoecklin, G. (2014). An optimized streptavidin-binding RNA aptamer for purification of ribonucleoprotein complexes identifies novel ARE-binding proteins. Nucleic Acids Res. 42. 10.1093/nar/gkt956.

38. Kreft, I.C., Huisman, E.J., Cnossen, M.H., Van Alphen, F.P.J., Van Der Zwaan, C., Van Leeuwen, K., Van Spaendonk, R., Porcelijn, L., Veen, C.S.B., Van Den Biggelaar, M., et al. (2023). Proteomic landscapes of inherited platelet disorders with different etiologies. J. Thromb. Haemost. 21, 359–372.e3. 10.1016/j.jtha.2022.11.021.

39. Silver, L.M., and Remis, D. (1987). Five of the nine genetically defined regions of mouse *t* haplotypes are involved in transmission ratio distortion. Genet. Res. 49, 51–56. 10.1017/S0016672300026720.

40. Mayya, V., Lundgren, D.H., Hwang, S.-I., Rezaul, K., Wu, L., Eng, J.K., Rodionov, V., and Han, D.K. (2009). Quantitative Phosphoproteomic Analysis of T Cell Receptor Signaling Reveals System-Wide Modulation of Protein-Protein Interactions. Sci. Signal. 84. 10.1126/scisignal.2000007.

41. Pollizzi, K.N., and Powell, J.D. (2015). Regulation of T cells by mTOR: The known knowns and the known unknowns. Trends Immunol. 36, 13–20. 10.1016/j.it.2014.11.005.

42. Aoki, K., Adachi, S., Homoto, M., Kusano, H., Koike, K., and Natsume, T. (2013). LARP1 specifically recognizes the 3′ terminus of poly(A) mRNA. FEBS Lett. 587, 2173–2178. 10.1016/j.febslet.2013.05.035.

43. Lahr, R.M., Mack, S.M., Héroux, A., Blagden, S.P., Bousquet-Antonelli, C., Deragon, J.M., and Berman, A.J. (2015). The La-related protein 1-specific domain repurposes HEAT-like repeats to directly bind a 5’TOP sequence. Nucleic Acids Res. 43, 8077–8088. 10.1093/nar/gkv748.

44. Fonseca, B.D., Zakaria, C., Jia, J.-J., Graber, T.E., Svitkin, Y., Tahmasebi, S., Healy, D., Hoang, H.-D., Jensen, J.M., Diao, I.T., et al. (2015). La-related Protein 1 (LARP1) Represses Terminal Oligopyrimidine (TOP) mRNA Translation Downstream of mTOR Complex 1 (mTORC1). J. Biol. Chem. 290, 15996–16020. 10.1074/jbc.M114.621730.

45. Herranz, N., Gallage, S., Mellone, M., Wuestefeld, T., Klotz, S., Hanley, C.J., Raguz, S., Acosta, J.C., Innes, A.J., Banito, A., et al. (2015). mTOR regulates MAPKAPK2 translation to control the senescence-associated secretory phenotype. Nat. Cell Biol. 17, 1205–1217. 10.1038/ncb3225.

46. Calo, E., Flynn, R.A., Martin, L., Spitale, R.C., Chang, H.Y., and Wysocka, J. (2015). RNA helicase DDX21 coordinates transcription and ribosomal RNA processing. Nature 518, 249–253. 10.1038/nature13923.

47. Cook, M.E., Bradstreet, T.R., Webber, A.M., Kim, J., Santeford, A., Harris, K.M., Murphy, M.K., Tran, J., Abdalla, N.M., Schwarzkopf, E.A., et al. (2022). The ZFP36 family of RNA binding proteins regulates homeostatic and autoreactive T cell responses. Sci. Immunol. 7, eabo0981. 10.1126/sciimmunol.abo0981.

48. Petkau, G., Mitchell, T.J., Chakraborty, K., Bell, S.E., D’Angeli, V., Matheson, L., Turner, D.J., Saveliev, A., Gizlenci, O., Salerno, F., et al. (2022). The timing of differentiation and potency of CD8 effector function is set by RNA binding proteins. Nat. Commun. 13, 2274. 10.1038/s41467-022-29979-x.

49. Zandhuis, N.D., Guislain, A., Popalzij, A., Engels, S., Popović, B., Turner, M., and Wolkers, M.C. (2024). Regulation of IFN-γ production by ZFP36L2 in T cells is time-dependent. Eur. J. Immunol., 2451018. 10.1002/eji.202451018.

50. Sandberg, R., Neilson, J.R., Sarma, A., Sharp, P.A., and Burge, C.B. (2008). Proliferating cells express mRNAs with shortened 3′ untranslated regions and fewer microRNA target sites. Science 320, 1643–1647. 10.1126/science.1155390.

51. Gruber, A.R., Martin, G., Müller, P., Schmidt, A., Gruber, A.J., Gumienny, R., Mittal, N., Jayachandran, R., Pieters, J., Keller, W., et al. (2014). Global 3′ UTR shortening has a limited effect on protein abundance in proliferating T cells. Nat. Commun. 5, 1–10. 10.1038/ncomms6465.

52. Nicolet, B.P., Jurgens, A.P., Bresser, K., Guislain, A., Bradariç, A., and Wolkers, M.C. (2024). Learning the sequence code of protein expression in human immune cells. Preprint, 10.1101/2023.09.01.555843.

53. Piccirillo, C.A., Bjur, E., Topisirovic, I., Sonenberg, N., and Larsson, O. (2014). Translational control of immune responses: from transcripts to translatomes. Nat. Immunol. 15, 503–511. 10.1038/ni.2891.

54. Rieckmann, J.C., Geiger, R., Hornburg, D., Wolf, T., Kveler, K., Jarrossay, D., Sallusto, F., Shen-Orr, S.S., Lanzavecchia, A., Mann, M., et al. (2017). Social network architecture of human immune cells unveiled by quantitative proteomics. Nat. Immunol. 18, 583–593. 10.1038/ni.3693.

55. Perez-Riverol, Y., Bai, J., Bandla, C., García-Seisdedos, D., Hewapathirana, S., Kamatchinathan, S., Kundu, D.J., Prakash, A., Frericks-Zipper, A., Eisenacher, M., et al. (2022). The PRIDE database resources in 2022: a hub for mass spectrometry-based proteomics evidences. Nucleic Acids Res. 50, D543–D552. 10.1093/nar/gkab1038.

56. Nicolet, B.P., Guislain, A., Van Alphen, F.P.J., Gomez-Eerland, R., Schumacher, T.N.M., Van Den Biggelaar, M., and Wolkers, M.C. (2020). CD29 identifies IFN-γ–producing human CD8 ^+^ T cells with an increased cytotoxic potential. Proc. Natl. Acad. Sci. 117, 6686–6696. 10.1073/pnas.1913940117.

