## Supplemental Figures for "mTOR signaling promotes cytokine production in T cells through 3’UTR-mediated translation control"

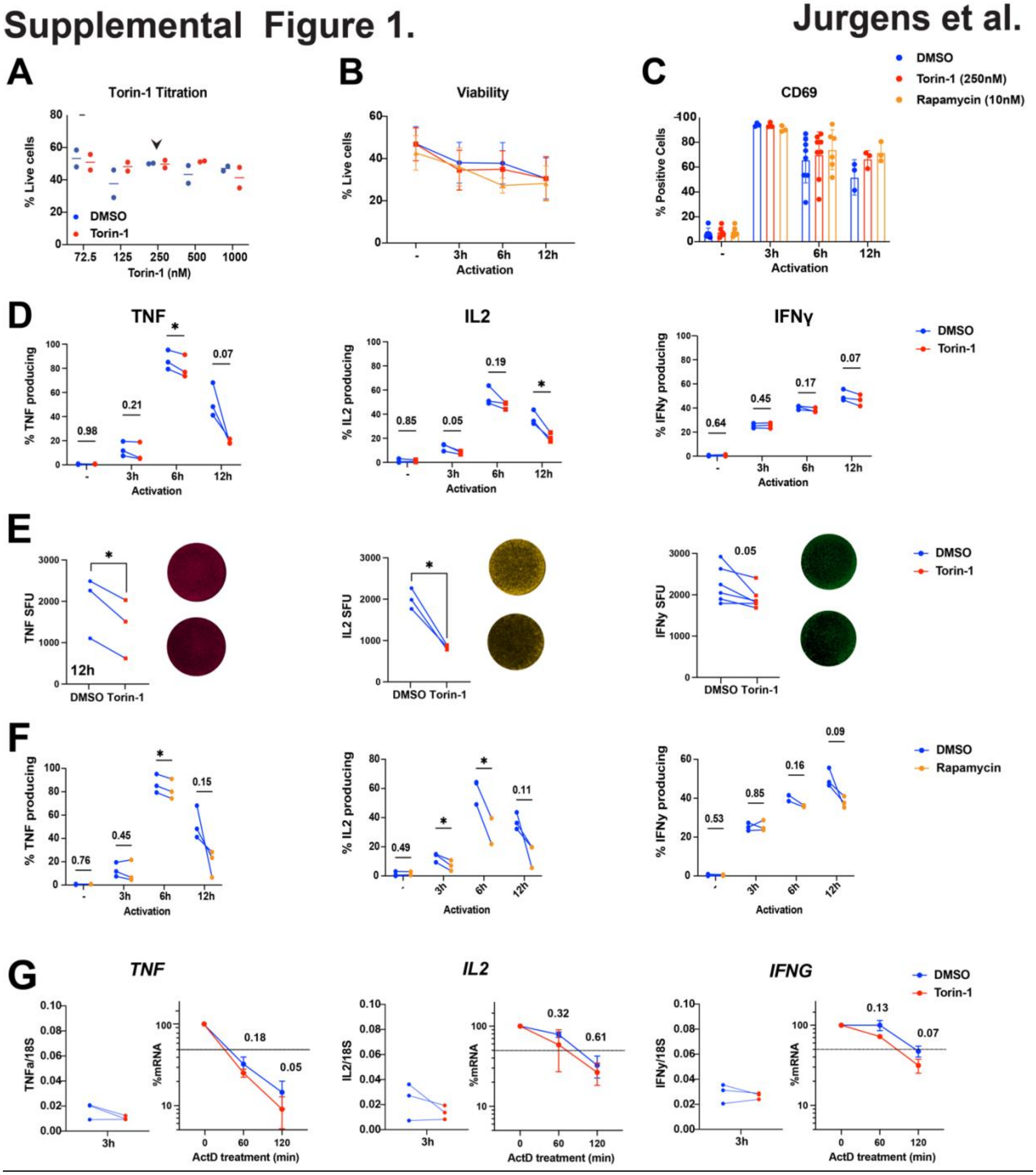


**Supplemental Figure 1. mTOR affects cytokine production in human CD8⁺ effector T cells**

A) Human CD8⁺ T cells were activated for 6h with PMA/Ionomycin in the presence of indicated concentrations of Torin-1 (red) or DMSO (blue). Arrow indicates concentration used in consecutive experiments.

B) -C) $CD8^+$ T cells were activated for indicated time points with PMA/Ionomycin in the presence of Torin-1 (red), Rapamycin (orange), or DMSO control (blue) and (B) measured for viability using near-IR dye, and (C) expression of the activation marker CD69. (-) indicates non-activated T cells that were treated with Torin-1, Rapamcyin or DMSO control for 12h. (n=3-6 donors, B-C: mean ± SD)

D) Indicated cytokine by intracellular cytokine staining (ICCS). Brefeldin A was added for the last 2h of activation. (n=3 donors, mean ± SD)

E) $CD8^+$ T cells were activated for 12h with αCD3/αCD28 in the presence of Torin-1 or DMSO control, and TNF, IL2 and IFNγ protein levels were measured by FluoroSpot. Left: representative FluoroSpot image of DMSO (top) and Torin-1 (bottom) treated cell. SFU: Spot Forming Units. SPU of $1x10^4$ cells plated per well. (n=3-6 donors, mean ± SD)

F) $CD8^+$ T cells were activated with PMA/Ionomycin in the presence of Rapamycin or DMSO control. Brefeldin A was added for the last 2h of activation. TNF, IL2 and IFNγ protein expression was measured by ICCS. (n=2-3 donors, mean ± SD)

G) $CD3^+$ T cells were activated for 3h with αCD3/αCD28 in the presence of Torin-1 or DMSO control. Left: Cytokine mRNA levels, normalized to 18S control. Right: T cells were treated for an additional 1h and 2h with actinomycin D (ActD), and mRNA levels were determined by RT-PCR. mRNA levels at 3h of activation was used as 100%. (n=3 donors, mean ± SD) B-G: one sided paired student's t test; $*p < 0.05$.

### Supplemental Figure 2 Jurgens et al.

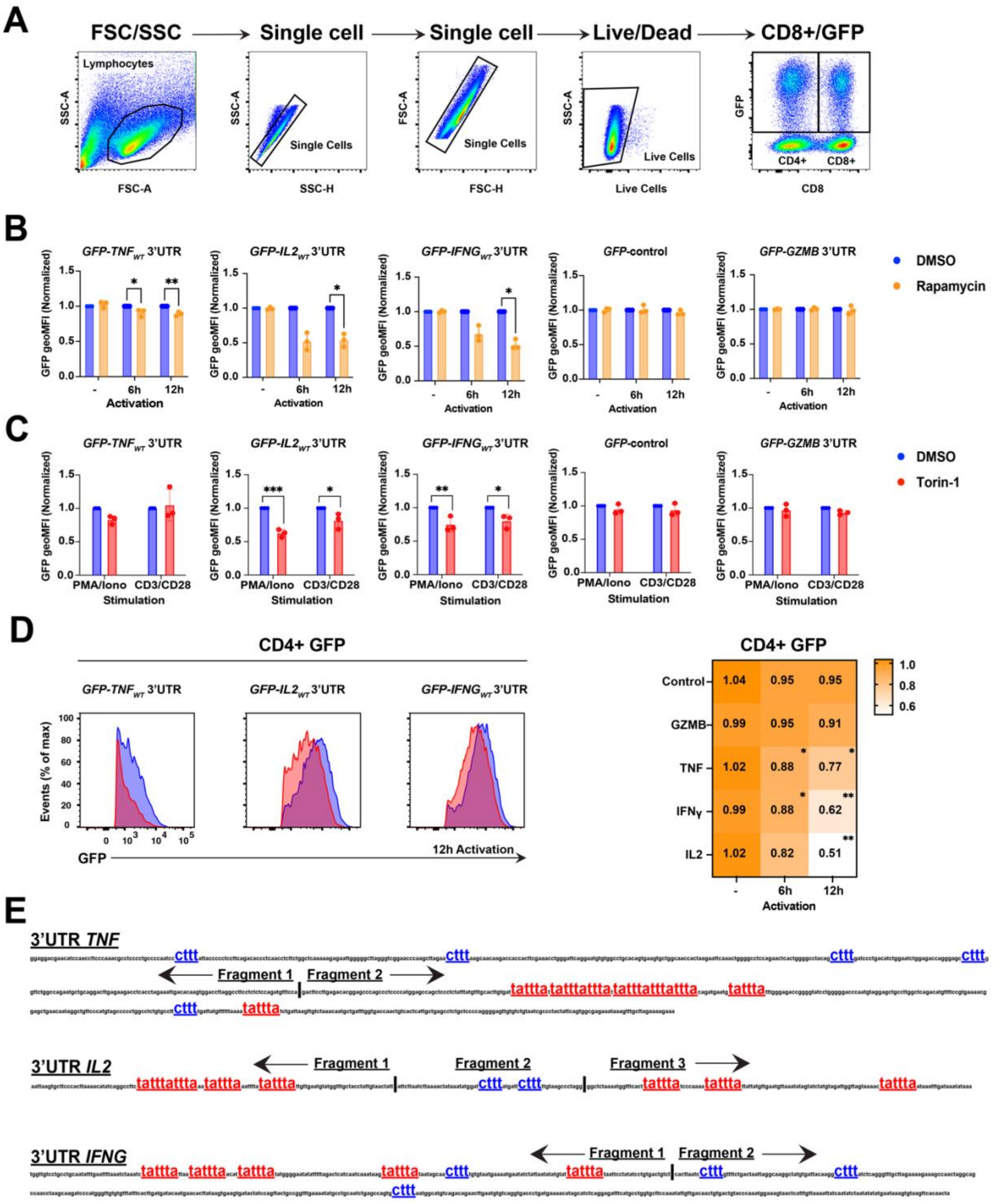


**Supplemental Figure 2. mTOR regulates cytokine production through the 3'UTR.**

A) Gating strategy of T cells expressing the GFP 3'UTR reporter constructs.

B) CD8$^{+}$ T cells expressing indicated GFP reporter constructs, or GFP *Control* were activated with PMA/Ionomycin in the presence of Rapamycin (orange) or DMSO control (blue).

(-) indicates non-activated T cells treated with Torin-1 or DMSO for 12h. GFP geoMFI of Rapamycin treated samples were normalized to the paired DMSO controls. (n=3 donors, mean ± SD).

C) CD8$^+$ T cells expressing indicated GFP reporter constructs were activated with PMA/Ionomycin or $\alpha$CD3/$\alpha$CD28 in the presence of Torin-1 (red) or DMSO control (blue) for 16h. GFP geoMFI was normalized to the paired DMSO control. (n=3 donors, mean ± SD)

D) CD4$^+$ T cells expressing indicated GFP reporter constructs were activated with PMA/Ionomycin in the presence of Torin-1 or DMSO control for indicated time points. Left: representative GFP expression at 12h post activation. Right: heatmap of GFP geoMFI of indicated reporter constructs. GFP *Control* and *GZMB* served as control. Heatmap scale represents GFP geoMFI ratio between DMSO/Torin-1 treated cells. (n=3 donors) B-D: one sided paired student t test; *p < 0.05, ** < 0.01, ***< 0.001

E) Sequence of the 3'UTRs of human *TNF, IL2* and *IFNG*. TOP-like CTTT motifs are indicated in blue, AREs in red. Vertical line with arrows represents design of cytokine 3'UTR fragments.

#### Supplementary Figure 3 Jurgens et al.

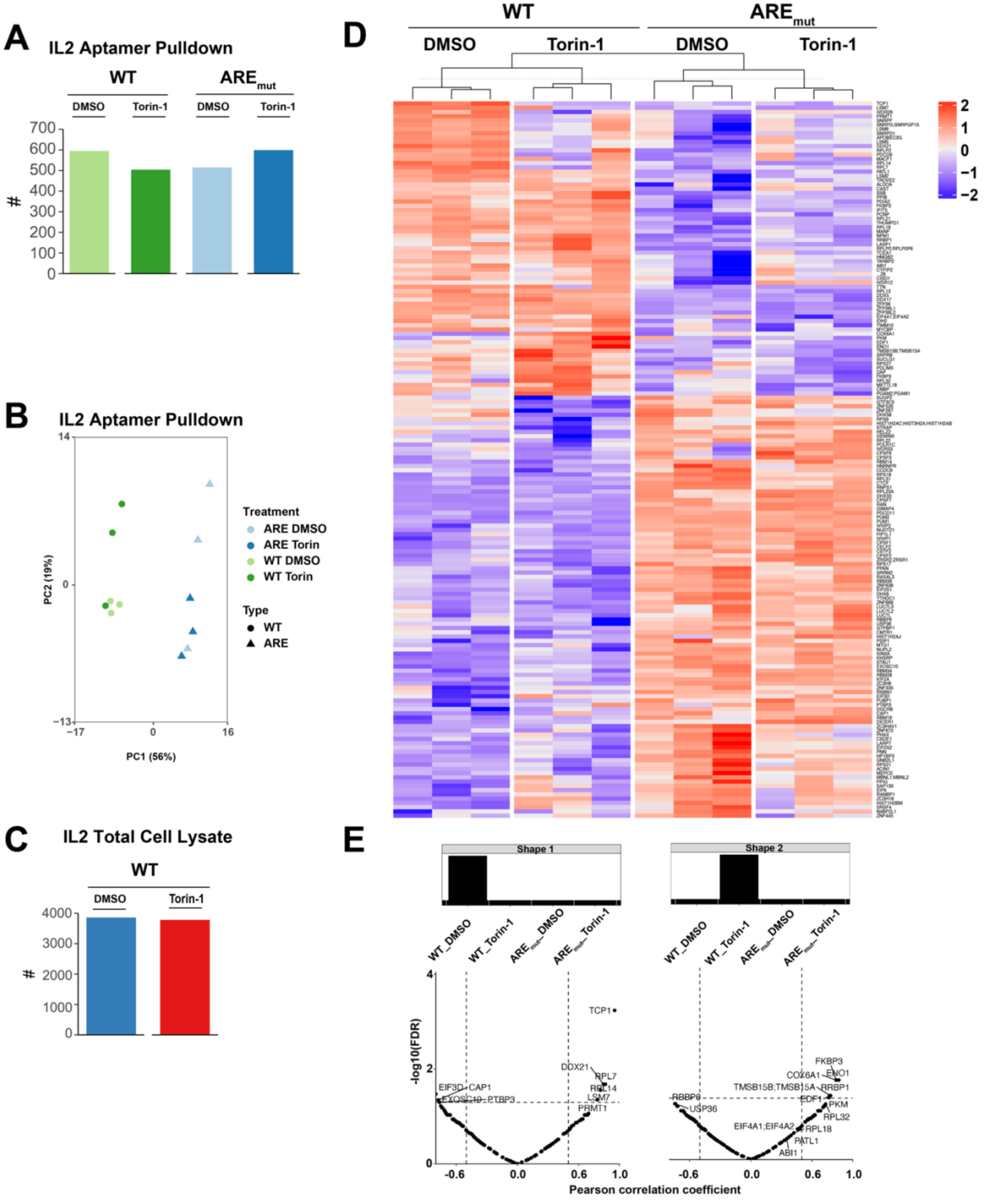

**Supplemental Figure 3. Identification of ARE-dependent and mTOR-mediated RBP binding to the *IL2* 3'UTR.**

A) Number of proteins found in *IL2* 3'UTR aptamer pulldown by Mass Spectrometry in different conditions as indicated. Bars are the mean of T cells from 3 donor pools consisting of 40 donors each. Each donor pool was activated for 3h with $\alpha$CD3/$\alpha$CD28 in the presence of Torin-1 or DMSO and total cell lysates were incubated with *in vitro* transcribed RNA as indicated (*IL2* WT 3'UTR vs *IL2* ARE-mut 3'UTR).

B) Principle component analysis (PCA) for indicated conditions.

C) Number of proteins identified in total cell lysates of A). Bars are mean of 3 donor pools.

D) Heatmap of all proteins (188) that were found enriched (LFC >1) in one of the four conditions in the aptamer pulldown. Color scale represents Z-scored log2 median-centered averaged intensities.

E) Shape analysis using supervised classification. Identification of proteins that were enriched in one condition, compared to all other conditions with a p<0.05 and LFC>1. Shape 1 represents proteins enriched only in *IL2* WT 3'UTR under DMSO conditions (ARE-Binding Proteins (ARE-BP) Torin repelled), and shape 2 represents proteins only enriched in *IL2* WT 3'UTR under Torin-1 conditions (ARE-BP Torin-1 induced). Correlation between hits and disorder specific theoretical protein profiles using a Pearson correlation coefficient of >0.5 (dotted line) and Benjamini-Hochberg adjusted P value <0.05 was used as threshold.

### Supplementary Figure 4 Jurgens et al.

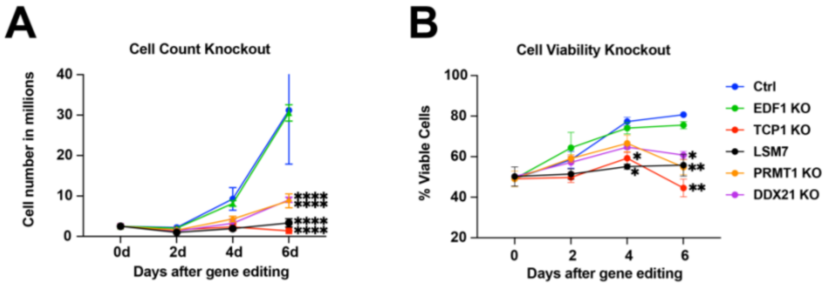


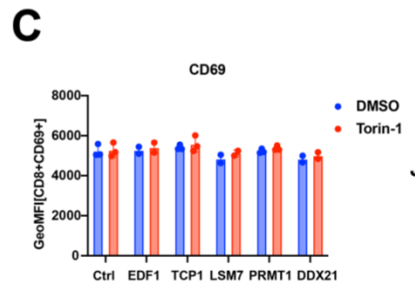


**D**

**DDX21 Phosphorylation status**

**Jurkat**

**Mayya *et al.* 2009 - OKT3 CD3 activation**

| Protein ID | Phosphopeptides | Phosphosites |
|---|---|---|
| DDX21_HUMAN | ---MPGKLRSDAGLESDTAM | 7 |
| DDX21_HUMAN | GKLRSDAGLESDTAMKKGETL | 13 |
| DDX21_HUMAN | LRSDAGLESDTAMKKGETLRK | 15 |
| DDX21_HUMAN | KAEPSEVDMNSPKSKKAKKKE | 71 |
| DDX21_HUMAN | TKKVTKNEEPSEEEIDAPKPK | 121 |
| DDX21_HUMAN | MRNGIDILVGTPGRIKDHIQN | 315 |

**CD4⁺ T cell (mouse)**

**Tan *et al.* 2017- 16h activation**

| Protein ID | Phosphopeptides | Phosphosites | Conserved | stim:no stim | log2(RAPTOR KO.16h/WT.16h) |
|---|---|---|---|---|---|
| Ddx21_murine | KQEEEPQDDTASTSK | 87 | No | 1.34 | -0.09 |
| Ddx21_murine | EIITEEPSEEEADMPKPK | 118 | No | 1.25 | -0.58 |
| Ddx21_murine | NGLSQPSEEEADIPKPK | 155 | Yes | 2 | -0.95 |
| Ddx21_murine | NGLSQPSEEEVDIPKPK | 189 | No | 2.53 | -1.24 |
| Ddx21_murine | NGLSQPSEEEVDIPKPK | 192 | No | 0.69 | -0.46 |
| Ddx21_murine | SNSSDAPGEESSSETEK | 235 | No | 2.69 | -0.8 |
| Ddx21_murine | SNSSDAPGEESSSETEK | 236 | Yes | 2.20 | -0.74 |
| Ddx21_murine | SNSSDAPGEESSSETEK | 243,244 | Yes | 1.74 | -0.65 |

**Supplemental Figure 4. mTOR regulates protein expression through the ARE-binding protein DDX21.**

A) Cell count and

B) % cell viability of RBP knockout $CD3^+$ T cells or control $CD3^+$ T cells with non-targeting crRNA (Ctrl) as defined by cell size by CasyCounter over the course of 6 days post CRISPR-editing. (n=2-4 donors, mean ± SD)

C) $CD8^+$ T cells expressing *GFP-IL2* 3’UTR were gene-edited for RBP knockout or non-targeting crRNA (Ctrl) using CRISPR-Cas9. Cells were activated with PMA/Ionomycin in the presence of Torin-1 (red) or DMSO (blue). C) geoMFI of CD69+ activation marker expression at 12h after activation. (n=3 donors, mean ± SD).

D) DDX21 phosphopeptides that were extracted from phosphoproteomics datasets of activated Jurkat cells (Mayya et al. 2009) and mouse T cells (Tan et al. 2017). Tan et al. data also indicate the log2 difference between Raptor (mTORC1)-deficient and WT cells. Red letter corresponds to phospho-site. A-B: One-way ANOVA with Tukey multiple comparison correction was performed. C-D: One sided paired student t test; (*p < 0.05, **p < 0.01, ***p < 0.001****p < 0.0001).
